# Decentralized Control of Insect Walking – a simple neural network explains a wide range of behavioral and neurophysiological results

**DOI:** 10.1101/695189

**Authors:** Malte Schilling, Holk Cruse

## Abstract

Control of walking with six or more legs in an unpredictable environment is a challenging task, as many degrees of freedom have to be coordinated. Generally, solutions are proposed that rely on (sensory-modulated) CPGs, mainly based on data from neurophysiological studies. Here, we are introducing a sensor based controller operating on artificial neurons, being applied to a (simulated) hexapod robot with a morphology adapted to *Carausius morosus*. We show that such a decentralized solution leads to adaptive behavior when facing uncertain environments which we demonstrate for a large range of behaviors – slow and fast walking, forward and backward walking, negotiation of curves and walking on a treadmill with various treatment of individual legs. This approach can as well account for these neurophysiological results without relying on explicit CPG-like structures, but can be complemented with these for very fast walking.

## Introduction

Motor control in animals deals with the control of a very high number of muscles (or degrees of freedom) that have to be controlled in a coordinated way in order to achieve complex behaviors. One key principle that allows animals to deal with such a complex control problem is modularization (D’Avella et al. 2015; Flash & Hochner, 2005; Mussa-Ivaldi 1999; Botvinick 2008), i.e. breaking the overall complexity of the control system down on a structural level (Binder et al. 2009). Research on locomotion in particular has highlighted decentralization as an organizing principle of such a modular structure that leads to parallel, local processing which is highly efficient, but requires a form of coordination.

Locomotion is a basic behavior of animals, the most important types of which are swimming, flying and walking, behaviors preponderantly characterized by rhythmic movement (Orlovsky et al. 1999). Locomotion has been studied in insects for a long time as this offers insights on different levels of analysis that are assumed to translate to other animals as well (Dickinson et al. 2000; Pearson 1995). On the one end of the spectrum, insects allow studying neuronal activities on a detailed neurophysiological level (Borgmann and Büschges 2015; Sponberg and Full 2008; Bender et al. 2010; Stein and Schmitz 1999). On the other hand, insect behavior can be observed in specific and well-defined contexts (Dürr et al. 2004; Ritzmann and Zill 2013; Cruse 1990; Bender et al. 2011). As a result, decentralization has been established as one key organizing principle, but there is disagreement on the internal organization of the local control modules and in particular if locomotion is driven by an intrinsic oscillation as such or by sensory input signals (Ijspeert 2008; Pearson et al. 2006; Borgmann and Büschges 2015; Ayali et al. 2015; Full and Tu 1991; Spence et al. 2010; Ritzmann and Zill 2013; Schilling et al. 2013a,b). Rhythmic biological movements – as long as these cannot be explained by mechanical, e.g., pendulum-like, properties – are generally considered to be controlled by intrinsic oscillations that are driven by neuronal systems and provide the basis of the required motor output (e.g., Daun-Gruhn and Büschges 2011; Ijspeert 2008, 2014; Orlovsky et al. 1999), a view strongly supported by studies on various types of animals. But most of the cases studied concern swimming or flying or fast running. The situation seems to be less clear when it comes to a different type of locomotion, namely walking. What might be crucial conditions that make a difference between (slow) walking on the one hand and swimming or flying on the other? The main difference appears to be given by the different environmental situations animals have to deal with (Bender et al. 2011; Borgmann and Büschges 2015).

Swimming and flying is operating in a fairly homogenous, kind of soft medium. Walking, in contrast, has to deal with a more difficult substrate providing unpredictable situations which may vary dramatically from step to step. Different to swimming and flying, walking is characterized by the necessity to orchestrate the movement of many joints. To cope with this problem, evolution has designed legs characterized by specific morphological properties (Bässler 1983). Each leg has several joints, at least three, in insects often four, in crustacea even more, leading to a high number of degrees of freedom to be controlled. As a result, at least 18 degrees of freedom (DoF), three per leg, exist in the case of a hexapod. 12 of these represent redundant degrees of freedom. This high number means that the kinematic system is characterized by “superfluous”, or redundant DoFs, which allow for highly adaptive behavior, but at the same time complicate the control problem (Bernstein 1967). The controller of such an underdetermined system is further challenged by the fact that these joints are not completely free to move, but are constrained by mechanical limitations among body and substrate making the control task even more complex (Schmitz et al. 2008).

In the remainder of this article, we will concentrate on walking of arthropods, with a focus on hexapods and will, first, start from a behavioral level of analysis, highlighting that walking comprises a behavioral spectrum with respect to different velocities that pose quite different challenges to the system. Later in the Introduction, we will turn towards related neurophysiological research that raises the question as to how intrinsic oscillatory systems may contribute to control of walking. The goal of this work is to provide a novel control model that for the first time attempts to cover the full range of these data.

### A continuum of walking behavior

Walking spans a whole behavioral continuum (Hoinville et al. 2015; Dickinson et al. 2000) – from slow walking to running. The contexts of the ends of this spectrum differ quite a lot. On the one hand, a typical and well studied example for slow walking can be found in the stick insect (Graham 1972; Borgmann and Büschges 2015; Dallmann et al. 2017; Bläsing and Cruse 2004). Stick insects normally climb through bushes and try to hide in twigs by mimicking them. They are moving slowly – less than three steps/s – as required by their behavioral context. When climbing through twigs there is no continuous substrate for walking. Instead, an insect has to search for footholds. During walking, antennae (and front legs to a certain degree) perform searching movements to sample the environment for footholds (Dürr 2001; Dürr 2014; Dürr and Schilling 2018). Furthermore, once a foothold has been found this information is shared with other legs that later-on will target and reuse that same foothold again (Cruse 1990; Dürr et al. 2004). In this case, the environment is dictating the conditions where to place a foot and in this way also the length of individual steps, which means that the stepping pattern may be quite variable. Temporal stability concerning the stepping pattern is less important in slow walking because longer stance duration allows for higher variability (Schilling et al. 2013a). Questions concerning postural stability are even less critical during slow walking as the insects can grasp the substrate with their tarsi necessary for climbing. To summarize, slow walking behavior is characterized by quite irregular coordination patterns as a consequence of the constant adaptation required by the given context.

At the other end, there is a large body of research on very fast walking in insects (Delcomyn 1971; Fuchs et al. 2011; Spence et al. 2010; Bender et al. 2011). Here, the situation differs considerably: For example, in cockroaches step frequency can be very high, up to 12-15 steps per second in *Blaberus discoidalis* (Sponberg and Full 2008; Weihmann et al. 2017; Bender et al. 2011); for *Periplaneta americana* even 25 step/s have been reported (Delcomyn 1971). For such very fast movements the delay of sensory pathways does play a role. Sensory information flow appears too slow to drive behavior selection (Sponberg and Full 2008). Instead, the control principle has to rely on an (in insects probably implicit) estimate of the continuation of the movement. The behavior is driven intrinsically which is supported by properties of the body (Holmes et al. 2006; Koditschek et al. 2004; Chiel et al. 2009): in running, postural stability becomes more a question of dynamic stability. Elasticities in the legs can support and stabilize the behavior as they are, for example, mechanically counteracting small disturbances in the substrate (Full et al. 2002). To summarize, fast walking is characterized as a rhythmic activity that leads to quite stereotypical coordination patterns of the legs being supported and stabilized by properties of the body.

Stressing these differences might lead to the – unintended – assumption that we are merely looking at different phenomena – different behaviors and different underlying control schemes. However, this is not necessarily the case. Fast walks are generally characterized by using a tripod pattern (i.e. at least three legs support the body at any moment of time). But tripod patterns can be observed in “slow” stick insects, too. In fact, there is a continuum (Graham 1972; Schilling 2013a) ranging from very slow walking (“pentapod” pattern sometimes called “wave gait”, at least five legs are on the ground at any time) over “tetrapod” pattern, sometimes called “metachronal gait” (at least four legs on the ground) to tripod pattern (at least three legs are on the ground). In cockroaches, *Periplaneta americana*, there is a tripod pattern from about a frequency of 5 steps/s up to an extreme speed of 25 steps/s (Delcomyn 1971; Ritzmann and Zill 2013). Below about 3 - 4 steps/s there is a continuous transition down to the slower tetrapod gait (Bender et al. 2011; Hughes 1952). In the stick insect *Carausius morosus* this transition occurs in a similar range (about 1 - 2 steps/s, Graham 1972). This suggests that there is a continuum of intermediate stable stepping patterns observed in probably all insects, at least in cockroaches and in stick insects, both representing the group of hemimetabolic insects, as well as in *Drosophila* (Wosnitza et al. 2013), representing a holometabolic insect. Recently, this view has been strongly supported by an excellent study of DeAngelis et al (2019).

Which stepping frequency might represent a limit above which sensory influence is not effective anymore? Sponberg and Full (2008) argue that for *Blaberus discoidalis* the fastest reflex response requires between 16 and 20 ms and assume that this delay is already too much to control switching between stance and swing states for a frequency of 10 steps/s (i.e. a period of 100 ms) or more. Results of Delcomyn (1991) with *Periplaneta americana* show that sensory feedback plays a role below 5 Hz but not anymore above this line. This means velocities up to 5 steps/s (period 200 ms) may be controlled by sensory feedback. Data of Zill and Moran (1981, *Periplaneta americana*) suggest a limit of 7 steps/s (for *Drosophila* this limit might be higher due to its small size and therefore shorter delays).

Here, we will focus on a controller structure that covers the full range of walking patterns from pentapod pattern to tripod pattern, but consider only patterns in a range that allows sensory inputs being effective. We will not yet deal with controllers that produce faster speeds and that are therefore assumed to underlie intrinsic oscillatory patterns (see however Discussion and Supplement).

### Free gait controllers: emergence of coordination patterns

We now want to turn towards a perspective that is focusing on underlying mechanisms. How may a controller be structured that is able to deal with the characteristics mentioned above? One assumption might be that specific behaviors observed may require specific neuronal modules (or motor schemas, see Cruse et al. 1990) which can be separately switched on or off (instructive examples are given by Steingrube et al. (2010) who distinguish eleven types of behaviors, seven of which are different walking types, or Daun-Gruhn and Tóth (2011) who use separate networks for tripod and tetrapod gaits).

However, as the behavioral data proposes that we deal with a continuous behavioral spectrum, this concept does not support a view of distinct motor schemas on the neuronal level (Graham (1972) for *Carausius;* Wosnitza et al. (2013) for *Drosophila;* see also Mantziaris et al. (2017) for discussion). Regular “gaits” only appear in the eye of the observer and it might therefore be misleading to assume that a controller has been evolved to produce separate gaits. Instead, it appears plausible that the controller has been designed to deal with general, unpredictable environments including unexpected disturbances, and that natural patterns strongly depend on specific environmental conditions (e.g. walking speed, or geometrical properties of the environment, for example when climbing over large gaps or along a vertical branch), where the “loop through the world” is assumed to play an important role (Brooks 1991). Similarly, straight forward walking or negotiating curves might not result from two separate modules (“straight forward” or “turning”), but may result from a system able to control turning by using various curvature radii (Dürr and Ebeling 2005; Rosano and Webb 2007; Gruhn et al. 2009; Cruse et al. 2009, Frantsevich and Cruse 2005) which includes straight forward walking. For further examples showing that not any behavioral phenomenon that we recognize requires a specific and explicit neuronal module, see Chittka and Neven (2009); Schmitz et al. (2008); Cruse and Wehner (2011); Hoinville and Wehner (2018) and Discussion.

Therefore, starting the design of a controller for the observed behavior by introduction of, at the outset, distinct structural elements might appear sensible in some cases, but should be avoided as long as no strong arguments support such a separation.

### Decentralization as a central control principle

To cope with the ability to walk, two problems have to be solved (e.g. Dürr et al. 2004). One concerns the question of how to control the movement of an individual leg and the other is how to couple the movement of the different legs. For both problems modularization appears to be a sensible solution (D’Avella et al. 2015; Hassabis et al., 2017).

The long tradition of behavioral research in insect walking has contributed largely to the understanding of how to couple the movement of different legs. Interleg coordination rules have been derived in diverse experimental settings that structure the overall behavior and coordinate the individual leg controllers. The whole spectrum of different gaits in a wide variety of contexts appear to emerge from such control structures. One example is given by our Walknet approach (Schilling et al. 2013a,b) in which coordination is based on individual leg controllers and on coordination influences regulating mainly the length of the step size. This system has been successfully applied in diverse contexts in dynamic simulation and on many robots.

Another intensively discussed question on this level of organization concerns how to control the behavior of an individual leg, i.e. how to select one of (at least) two behaviors: producing a stance movement in order to support and propel the body or a swing movement, during which the leg is lifted off the ground and moves into the opposite direction to start a new stance. As addressed above, the behavioral evidence may provide different answers depending on the context and the velocity of walking. On the one hand, slow walking requires integration of detailed feedback from the environment. On the other hand, very fast walking, or running, cannot keep up with integration of sensory input due to sensory delays. In this context, the controller has to produce a rhythm on its own. To cope with these different requirements, it is generally assumed that production of intrinsic rhythm represents the basic function and that such systems are, during slow walking, influenced by sensory feedback to allow adaptation to properties of the environment. Here we show that a slightly changed concept can explain more data by assuming that in the intact animal intrinsic rhythms are only applied during very fast walking. Our model shows that a form of mutual inhibition as a key organizing control principle is sufficient to explain, on the one hand, sensory switching of behaviors on the single leg level, and on the other hand, that more stable gaits and (seemingly) fixed coordination patterns emerge when the same structure is driven with a higher velocity. Nonetheless, this controller is able to deal with the full spectrum from pentapod to tripod walking, without relying on intrinsic rhythm generators. Beyond a given step frequency different connections may start to dominate (and in the end completely override) sensory feedback.

Both sides of the discussion that concerns either intrinsic rhythms or sensory-driven control often focus on quite specific experimental settings. Therefore, in order to show that the proposed control structure addresses the whole spectrum, it has to be tested in all of these contexts. Thus, in the next section we briefly summarize the view of intrinsic rhythms which largely puts a focus onto a different level, as are detailed neurophysiological experiments, not considered in Walknet.

### Central pattern generators

As mentioned above, intrinsic rhythms are often assumed to be caused by central pattern generators (CPG) (Ijspeert 2008; Orlovsky et al. 1999). The recognition of CPGs as the basic control system is based on experimental findings made by studies of deafferented animals (Pearson 1995; Orlovsky et al. 1999). Deafferentation is realized operationally by interrupting sensory input and motor output and later, in addition, by stimulating the neural system by, for example, application of pilocarpine (e.g. Büschges et al. 1995). As detailed below, such experiments could demonstrate that motor neurons driving antagonistic muscles of a given joint show – usually low frequency (in the order of 0.2 cycles/s) – oscillations with some similarity to those shown in intact moving animals. As these studies cover a broad range of species, both vertebrates and invertebrates, these results have led to the generally acknowledged assumption that such CPGs are responsible for controlling the basic rhythm observed in locomotion (Orlovsky et al. 1999). In addition, sensory feedback is assumed to be required for modulation, for example for resetting these CPGs allowing for adaptation to varying environmental conditions (Borgmann and Büschges 2015; Ritzmann and Zill 2013).

In the cases of swimming or flying, the temporal coordination between different CPGs observed in the deafferented animals shows in-phase coupling in particular between contralateral neighbors, which correspond relatively well to coupling found in the intact, behaving animal (see below). This is in contrast to the case of walking behavior that is characterized mostly by anti-phase coupling of neighboring legs (Graham 1972; Wosnitza et al. 2013; Delcomyn 1971). Here, the coordination of deafferented leg controllers is often quite different, eventually even opposite to that of normal walking (e.g. Büschges et al. 1995; Knebel et al. 2017; Borgmann et al. 2007, 2009; Mantziaris et al. 2017; we will deal with an interesting different case (Couzin-Fuchs et al. (2015); Johnston and Levine (2002) in the Discussion and Supplement).

This situation is highlighted by a detailed recent study on locusts (Knebel et al. 2017), focusing on interleg coordination in deafferented locusts (recordings from motor neurons activating the depressor muscle). The authors argued that the temporal patterns observed in the different thoracic ganglia may provide the basis for that found in walking animals. However, coordination in walking locusts is quite different to that found in the deafferented, in situ preparations. In general, in fully or partly deafferented insects ipislateral neighboring legs fire in-phase, whereas in walking these legs typically show about anti-phase coupling. Supporting results for *Carausius* are presented by Mantziaris et al. (2017), and Borgmann et al. (2007, 2009), who recorded from the motor neurons activating the retractor muscles, and by Büschges et al. (1995), who recorded from motor neurons of all three joints. Interestingly, corresponding results have been found for crustacea (Clarac and Chasserat 1979; Sillar et al. 1987). Furthermore, Knebel et al. (2017) found in-phase coupling also between contralateral legs if all three thoracic ganglia were treated with pilocarpine, which again contrasts to normal walking behavior, but can be observed in swimming (Ikeda and Wiersma 1964, crayfish swimmerets) and flying (Pearson 1995, locust). These results raise questions concerning the functional properties of these oscillatory systems.

### Simulations as a functional tool to analyze principles in a behavioral context

These questions cannot be approached by the current Walknet (Schilling et al 2013a,b). Thus, we strive for a solution that allows to maintain the basic structure and behavioral properties of Walknet, but at the same time is realized by an artificial neuronal network with neurons endowed with dynamic properties and that can be analyzed on a neural level. Furthermore, the motor system is organized using an antagonistic architecture (Szczecinski et al. (2013); Rubeo et al. 2018; Schumm and Cruse 2006; Cruse 1980; Daun-Gruhn and Tóth 2011) as such a more fine grained neuronal anatomy may allow for more detailed interpretations of the control structure proposed. We will not deal with an interesting series of studies on hexapod walking published by Manoonpong, Wörgötter and colleagues, based on Manoonpong et al. (2008), as our main focus is not to construct a hexapod robot being inspired by ideas taken from insect studies. Rather, our goal is to search for a simulation that allows to better understand specific biological systems, in particular, the stick insect *Carausius morosus* (see Fig. 1).

**Fig. 1.**
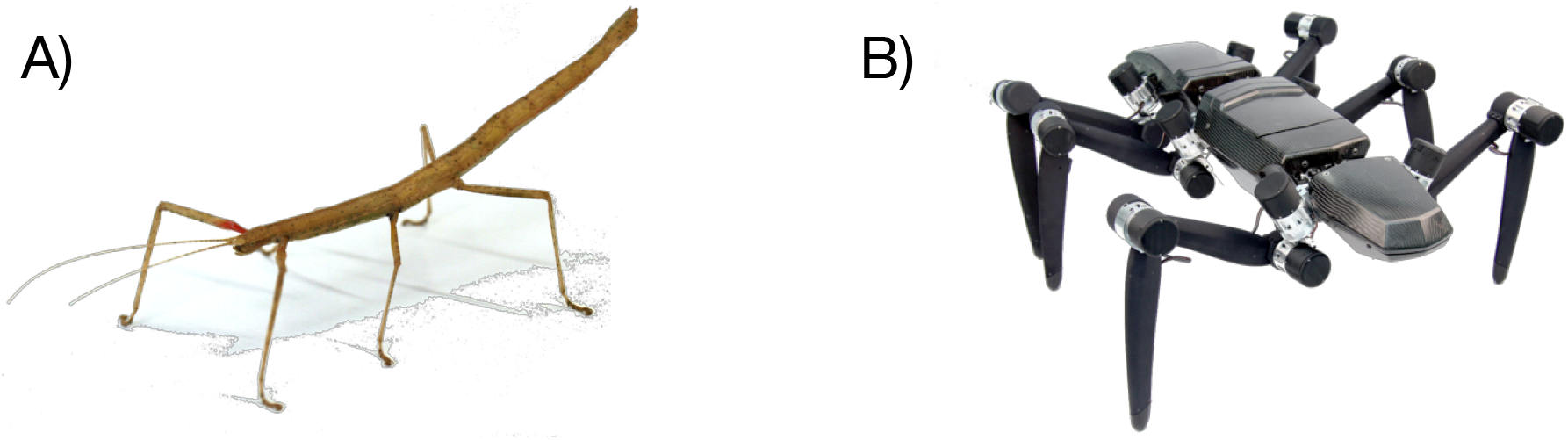
(a) Indian stick insect Carausius morosus. (b) Robot Hector which is used in simulation for testing the control principles derived.

For our approach, what is required further are a body, legs, muscles (motors) for each joint, sensors, and a physical environment. This property is important, because the existence of a body with legs coupled via the environment may allow simplifying computations (Brooks 1991). Concerning simulation of the body, software simulation and hardware simulation may be used. Here we start with a dynamic software simulation of the robot Hector (Schneider et al., 2014).

Following neurophysiological studies on insects (Mu and Ritzmann 2005; Tryba and Ritzmann 2000; Watson and Ritzmann 1998) each artificial motor neuron produces velocity signals to drive its muscle. Muscle properties are approximated by motors equipped with an elastic element. With such an architecture, the following general questions will be addressed.

i. How may (velocity dependent) interleg coordination (Dürr et al. 2004) be realized on the neuronal level in a way that temporal patterns will emerge that correspond to the patterns observed in behavior and that can adapt to external disturbances?
ii. How could the three joints of a single leg be coordinated? The joints of a leg are often not moved in-phase, but, depending on the current environmental situation, may show varying phase values. Negotiation of curves adds to the complexity of the task. Various solutions for intraleg coordination have been proposed (Daun-Gruhn and Tòth 2011; Büschges 2012; Rubeo et al. 2018; Steingrube et al. 2010), but have not been tested in the full range of experimental data available.
iii. Is the principle of mutual inhibition as applied in the proposed architecture sufficient to explain intrinsic oscillations that emerge in specific experimental situations (e.g., Knebel et al. 2017; Borgmann et al. 2009; Clarac and Chasserat 1979)?

Overall, the goal of the approach is not to reproduce specific, singular observations looked at in isolation, e.g. a specific gait type. In contrast, our goal is to establish a controller, a holistic model based on a minimum of a priori assumptions, that is able to describe a large amount of known behaviors that emerge from its decentralized structures (Schilling et al. 2013a). These behaviors concerns the full continuum of stepping patterns as observed in insects from pentapod via tetrapod to tripod patterns including intermediate stable patterns (Graham 1972; Wosnitza et al. 2013), backward walking (Graham and Epstein 1985; Jeck and Cruse 2007) and negotiation of curves (Dürr and Ebeling 2005; Rosano and Webb 2007; Gruhn et al. 2009) as well as stability against disturbances of leg movements studied in many experiments (for a review see Cruse 1990). Furthermore, a number of neurophysiological studies should be described that demonstrate the appearance of intrinsic oscillation also in slow walking arthropods, both insects and crustaceans (e.g., Knebel et al. 2017; Borgmann et al. 2009; Clarac and Chasserat 1979) which do, however, not contribute to normal, slow walking. Such a simulation may provide a holistic description, suited for testing hypotheses as well as to stimulate new experimental approaches.

## Results

The proposed neuronal architecture (for details see Material and Methods) has been tested in a variety of different experimental situations, ranging from diverse behaviors to analyses of neural activity. It follows the basic characteristic of Walknet (Schilling et al. 2013a,b), but is now constructed using a detailed neuronal architecture (Fig. 2 and Methods). Central to this control approach is the notion of decentralization realized as individual controllers for each leg. Front legs, middle legs and hind legs are controlled by a ganglion each, as found in the stick insect: the prothoracic ganglion, the mesothoracic ganglion and the metathoracic ganglion, respectively. Each ganglion consists of two hemiganglia, right and left, which control the corresponding legs. Interleg coordination is organized by local rules acting between neighboring legs. On the one hand, a set of these rules influences step length (coordination rule 1, 2 and 3, represented in Fig. 2 by brown units, see also Fig 11). On the other hand, as an additional rule that has been introduced into the Walknet control structure (Schilling et al. 2007) rule 5 (Fig 2, shown in ocher) is incorporating load-dependent signals into the controller.

**Fig. 2.**
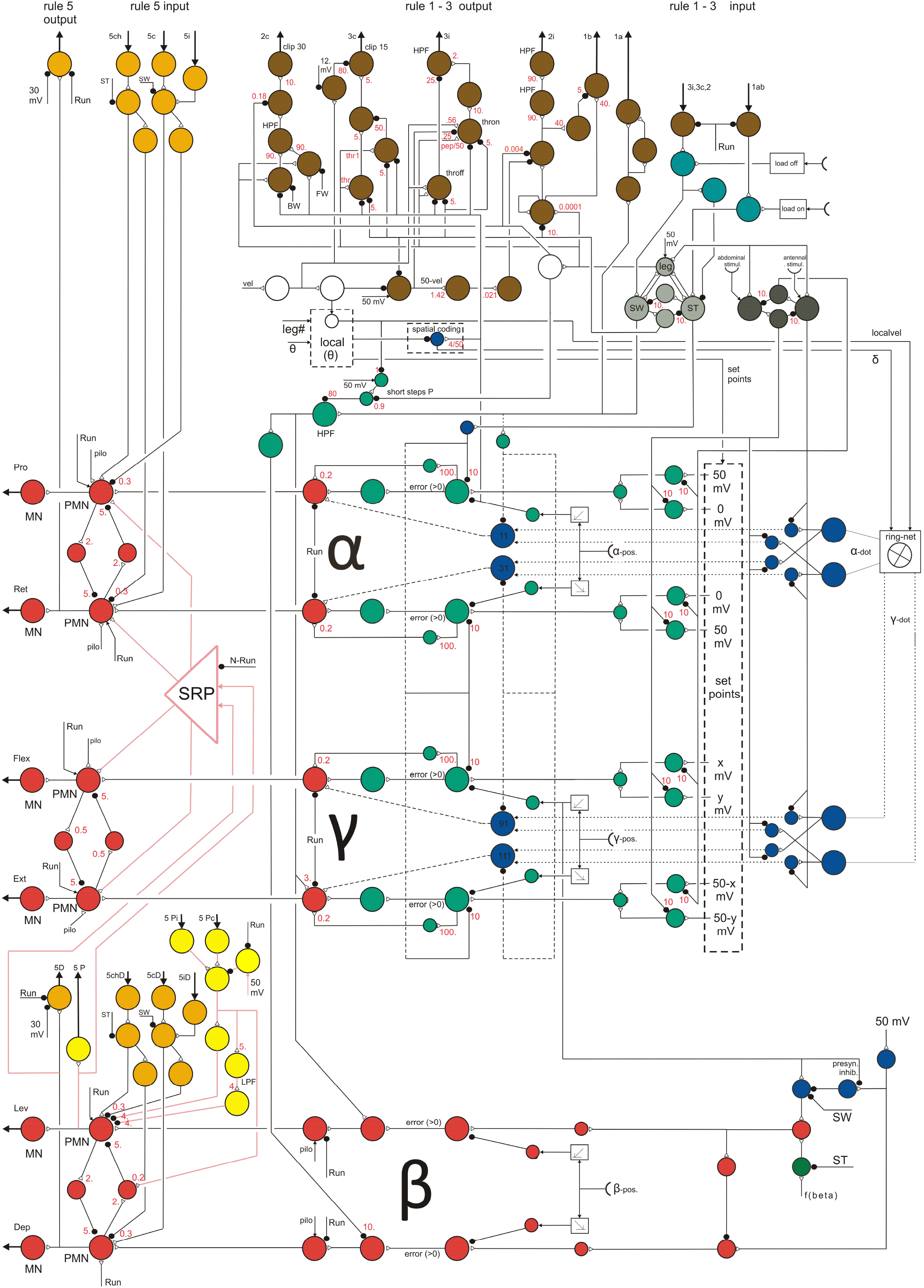
Controller for one leg. Artificial neurons are depicted as colored circles. Sensory input structure for swing: green, sensory input structure for stance: blue. Common output: red. Coordination channels: brown for rules 1 – 3, ocher for rule 5, light yellow for rule P (Pearson rule, see Supplement) with connections in pink (Suppl.). Motivation units: light grey (swing – stance), dark grey (forward – backward), white (central units). For details of ring net see Fig. 10. SRP net and its’ connections (pink) are detailed in Suppl. (Fig. S6). Joint alpha (α): Pro – protractor, Ret – retractor; joint beta (β): Lev – levator, Dep – depressor; joint gamma (γ): Flex – flexor, Ext – extensor. Black dots: inhibitory synapses, open black triangles: excitatory synapses. For detailed explanations see text and supplement.

The movement of a walking leg is characterized by two states, stance and swing. Following the antagonistic structure of the biological motor system the output of the individual joint muscles is controlled by an antagonistic structure, too. Each muscle is driven by a motor neuron (Fig 2, leftmost red units, MN). To minimize co-contraction of antagonistic muscles, premotor units (Fig 2, red, right neighbor of each motor unit, PMN) are connected via inhibitory units forming a four-unit recurrent network. The input to the controller (apart from local sensory feedback, Methods) is given by one unit (Fig 2, upper part, below the brown units representing coordination influences, leftmost white unit, input “vel”) that represents the desired global walking velocity (given in mV between 0 and 50 mV). The output of the controller, i.e., the motor neurons (MN), is a velocity signal that is used to drive the joint (for further explanation see Material and Methods). This controller architecture is assumed to encapsulate quite general control principles across different animal species (Dickinson et al. 2000).

In order to show the reach of these principles, the controller is applied and tested on a scaled up robotic model of an insect even though the dynamics in this scaled version are changing in a non-favorable way. As an insect, the model has three pairs of legs, front legs, middle legs and hind legs (FL, FR, ML, MR, HL, HR, with L for left and R for right leg). Each prototypical leg is equipped with three joints, termed alpha joint, beta joint and gamma joint. The alpha joint can be abstracted to be driven by two antagonistic muscles, protractor and retractor, essentially moving the leg from rear to front and back, the beta joint, with two antagonistic muscles levator and depressor, moving the leg up and down, and the gamma joint with two antagonistic muscles termed flexor and extensor, which essentially decrease or increase the length of the leg.

To illustrate the simulation results, apart from videos, so called foot fall patterns will be used: The temporal orchestration of the six legs are depicted by black bars showing the swing state (Figs 3, 4, 6, S3, S4). In other cases (legs being deafferented) black bars illustrate the activation of specific motor neurons when above a given threshold (Figures 7–9, S5, S7).

**Fig. 3 A-H.**
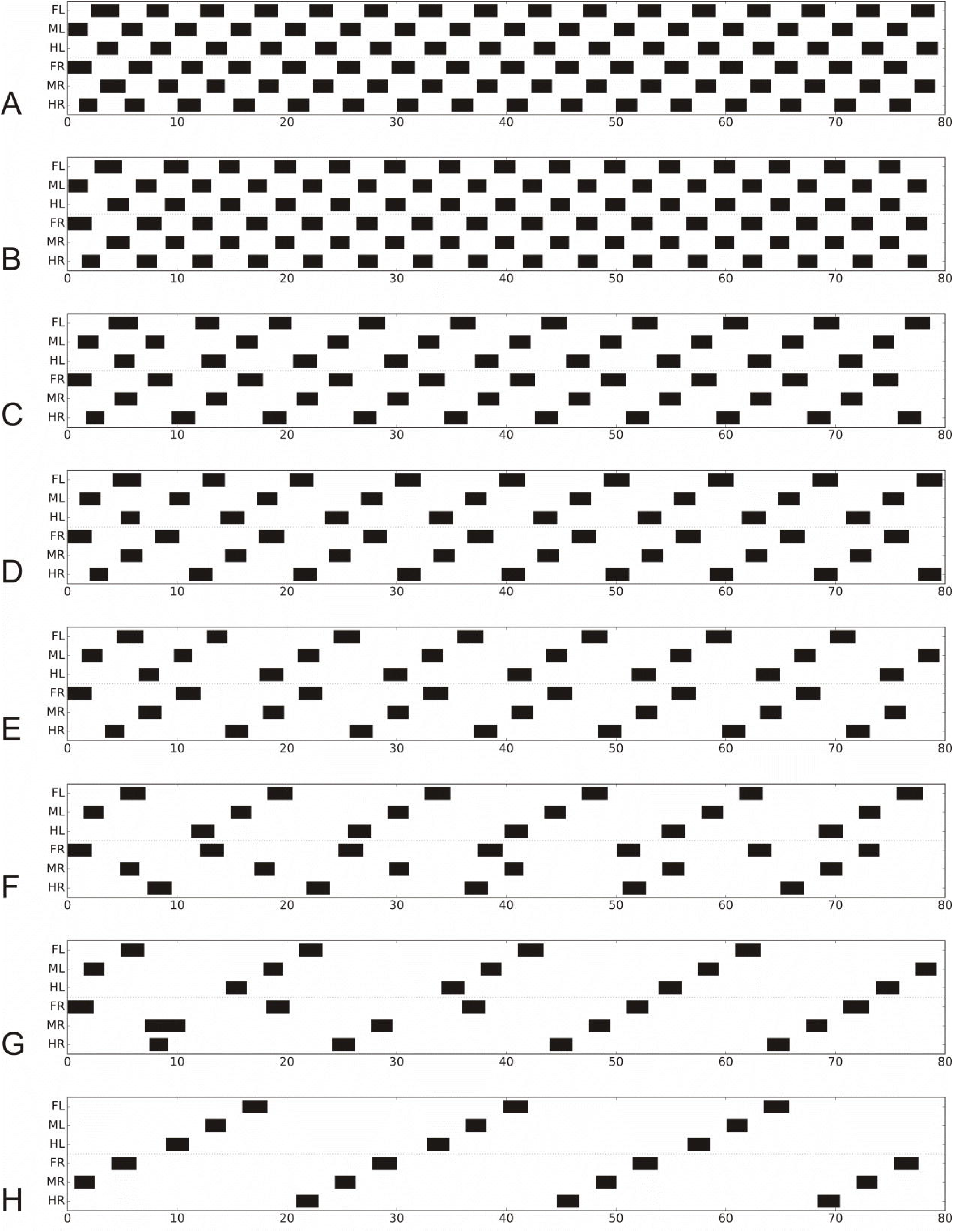
Footfall patterns for velocities driven by 50 mV, 40 mV, 30 mV, 25 mV, 20 mV, 15 mV 10 mV and 8 mV (G: the latter after disturbance of MR at about 8-11 s, see prolonged swing) and 8 mV (H: a stable pattern reached after more than 240 s), for undisturbed patterns see Fig S4 C, D). Apart from the last two runs all started with the same leg configuration. The corresponding transient phases for the last two runs are given in supplement (S4 C,D). Black bars show swing mode, Legs: FR front right, MR middle right, HR hind right, FL front left, ML middle left, HL hind left. Abscissa: Time (s).

**Fig. 4.**
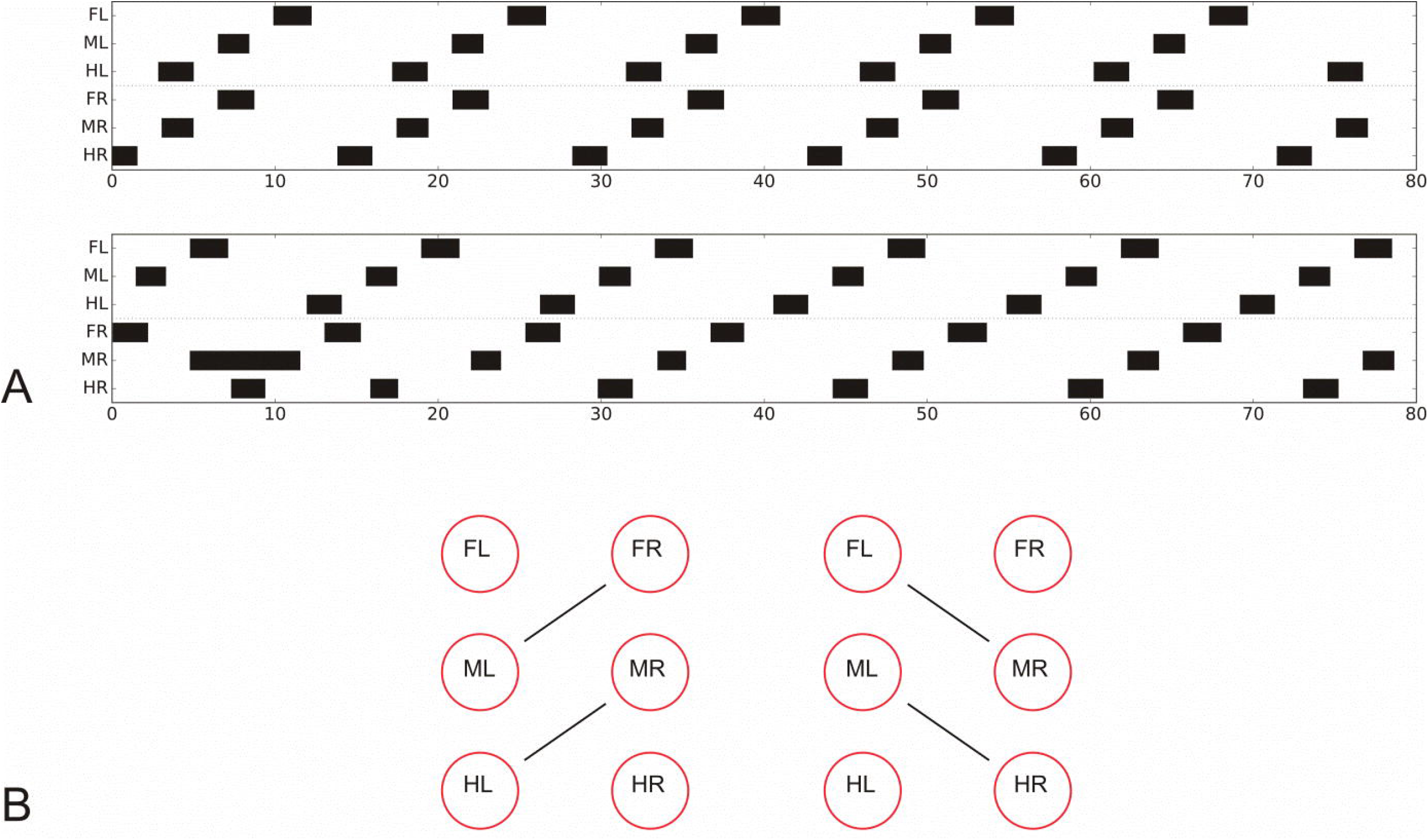
Footfall patterns for velocity 15 mV. Upper panel (A) as in Fig 3F, after a stable patter has been reached; lower panel (B) which, after a disturbance of the right middle leg during swing, represents a mirror image version. B) illustrates which leg pairs are coupled together in either case.

In the following, first, behavioral results will show that neuroWalknet can produce diverse walking behaviors: different gait patterns emerge from different velocities, the system can negotiate curves (shown in the supplement) and can deal with backward walking. Second, neural activities of the control structure will be analyzed in detail in order to compare these results with the findings of intrinsic oscillations as found in various neurophysiological experimental settings, for example dealing with deafferented animals.

### Forward walking – emerging gaits

Which patterns are produced when neuroWalknet is walking using all six legs and with different velocities? Fig 3 shows footfall patterns for different velocities, ranging from fast to very slow walking. Velocities are fed into the system as an external input and as a neural activation (Fig. 2, leftmost white unit). Here, inputs of 50 mV, 40 mV, 30 mV, 25 mV, 20 mV, 15 mV 10 mV and 8 mV were used (we also tested 45 mV and 35 mV, see Suppl. Fig. S4, for corresponding videos see Supplement). All walks are shown for 80 seconds. In the simulations shown, the robot always started from the same posture. As this was close to a tripod pattern, for higher velocities this posture already fitted the temporal pattern quite well, and accordingly a stable pattern was reached very soon for velocity signals induced by 50 mV and 40 mV. This starting configuration was however not optimally suited for most of the slower walks. Nonetheless, stable patterns emerged fast for velocities down to 20 mV and after only a couple of steps for 15 mV. In the other cases (vel 10 mV or 8 mV) more steps were required to reach a stable state. Fig. 3 (vel 10 mV) shows a case in which a stable pattern was reached quite fast for a slow velocity due to introduction of an additional disturbance (a deliberate prolongation of the first swing of the right middle leg). Fig. 3, vel 8 mV, shows a stable state reached after more than 240 s. The transition patterns found when using the generally applied starting configuration without additional disturbances are depicted in Supplement for 240 s each (Fig. S4).

Using the traditional characterization, patterns driven by velocities of 50 mV or 40 mV correspond to tripod gait, patterns produced between 30 mV and 15 mV are obvious tetrapod gaits, whereas patterns produced around 35 mV or between 15 mV and 10 mV (not shown) may be termed intermediate gaits, or perhaps tetrapod, depending on the exact definition. Patterns of 10 mV and below correspond to pentapod gaits. These results show that stable patterns emerged from the control structure and the interaction with the environment showing adaptive behaviors that can deal with disturbances given by uncomfortable starting configurations or deliberate manipulation (see Discussion).

Whereas typical tripod gait patterns and wave gait patterns are symmetrical with respect to right and left legs, this is not the case for the group labeled tetrapod. Here, for a given velocity, two mirror image versions are theoretically possible. Indeed, both versions can be produced when the walker is disturbed in some way (or, correspondingly, another starting configuration had been used). This is shown for vel = 15 (Fig 4), where Fig 4A, upper panel, shows the pattern directly following the undisturbed version as presented in Fig 3, whereas Fig 4B, lower panel, shows a walk with the same starting configuration, but a disturbance applied after about 15 s by artificially prolonging the swing duration of the right middle leg. Fig 4C illustrates which leg pairs swing together in either case. Note that these patterns show a contralateral phase different from 0.5, which agrees with experimental findings observed during tetrapod gait (*Carausius:* Graham, 1972; *Drosophila:* Wosnitza et al. 2013). For higher and for lower velocities, contralateral phase approaches values of 0.5. Note that averaging mirror image patterns with contralateral phase values of 0.33 and 0.66, for example, would lead to a mean phase value of 0.5.

To compare the footfall pattern received in the simulation with data obtained from stick insects (Graham 1972, his Fig 7), Fig 5 shows original data of Graham (his “3L1”, i.e. the time lag between beginning of swing in the (e.g. left) hind leg and the beginning of swing in the ipsilateral front leg vs. leg period). The corresponding values found in the simulation are depicted by red dots. Qualitatively, there is good agreement. Note that time scales differ by a factor of around 10 (this has been adapted as a consequence of the scaling of the robot structure).

**Fig. 5.**
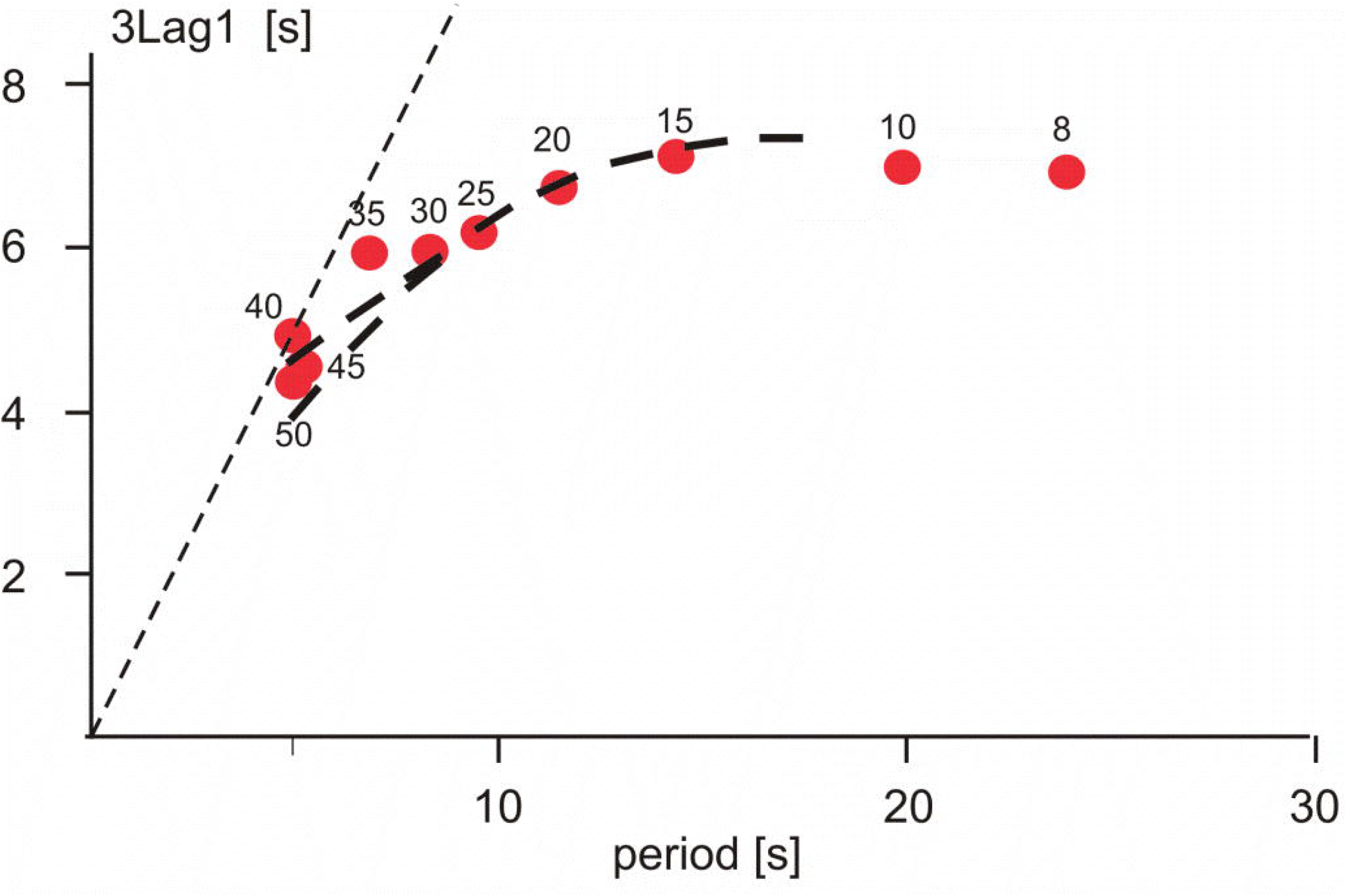
Duration of lag 3L1 (time lag between beginning of swing in hind leg and beginning of swing in the following front leg), vs. period (duration of swing plus stance) as proposed by Graham (1972). Time (s). Dashed lines show the average data from Graham (1972), his Fig 7; time *0.1)

Starting or stopping a walk is simply reached by setting the velocity unit from zero to the desired value, or back to zero (not shown). When the walker is stopped, legs that are in swing state finish swing movements, so that at the end all legs adopt ground contact as observed in insects. Controlling start and stop of a hexapod using a CPG-based controller (Tóth and Daun 2019) requires a considerable effort to tame the dynamic properties of the network. This is an interesting result as it highlights the problems CPG based controllers have to deal with when realistic behavioral tasks have to be solved (related problems are found by Rubeo et al 2018, too).

An example for negotiating curves is given in the Supplement (Fig S 3). Furthermore, in addition to the specific cases mentioned above, systematic disturbances have been performed by prolonging swing states during forward walking. In all cases studied, the controller reached a stable walking pattern (results not shown, see Discussion, and see also the first steps of backward walking (Fig 6).

**Fig. 6.**
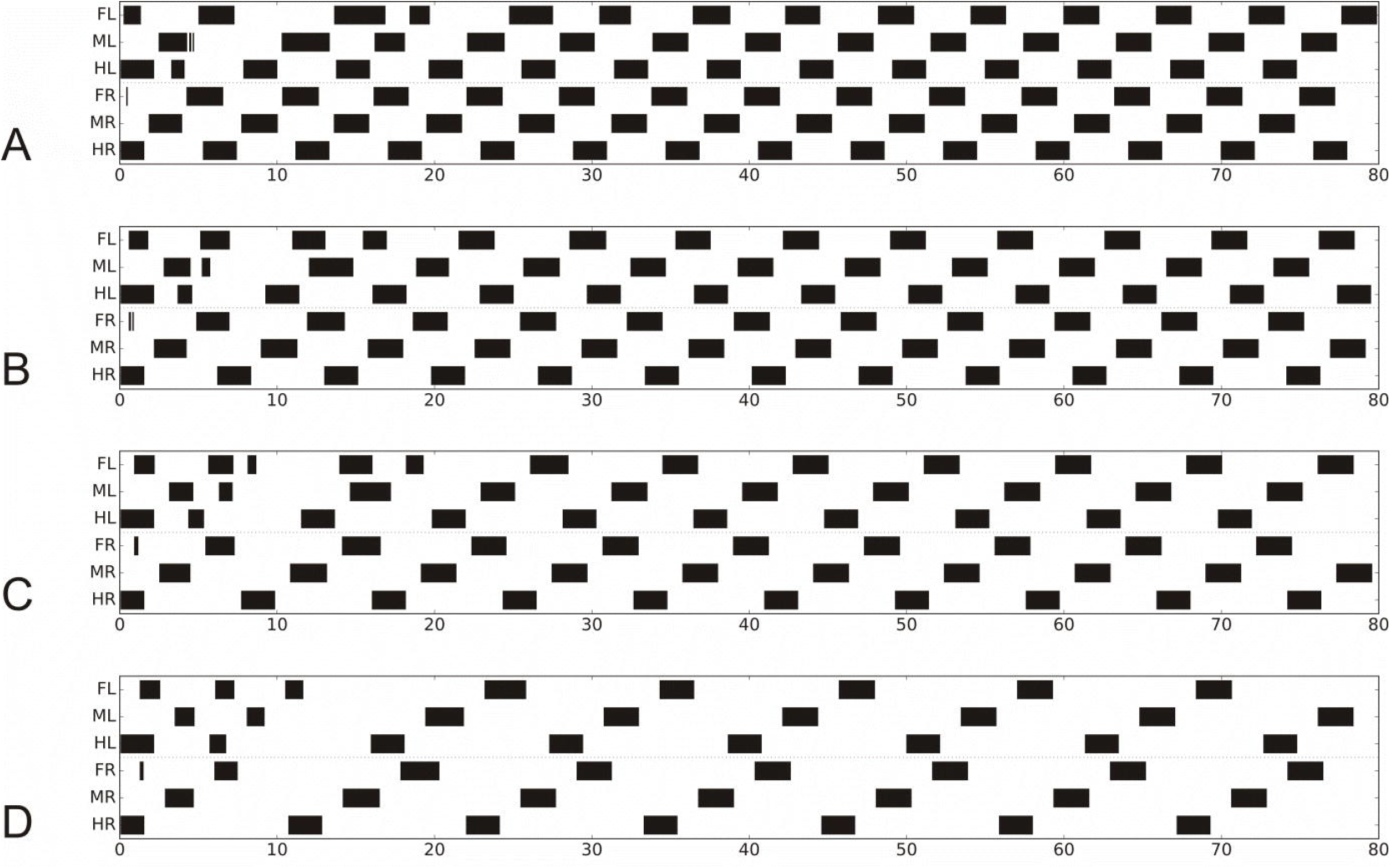
Footfall patterns, backward walking for velocities driven by 50 mV, 40 mV, 30 mV and 20 mV. Abscissa: time (s). For further details see Fig 3.

Taken together, the simulations provide a good qualitative description of the experimental data, showing typical “gait” patterns, allowing for symmetry breaks and qualitative agreement concerning the ipsilateral delay depending on velocity. Further, contralateral phases of about 0.5 are observed for fast and for very slow movements and deviating values (around 0.7, 0.3) for intermediate velocities. Furthermore, there are stable patterns that might not be characterized as tripod or tetrapod. Thus, using this control architecture, adaptive walking emerges with different patterns only depending on a selected velocity.

### Backward walking

Whereas forward walking and negotiation of curves form a continuum with the turning radius between zero and infinity, backward walking represents a separate state. A simple solution to introduce backward walking is to activate only rules 1 and 2i for ipsilateral coupling and use a version of rule 2c adapted to backward walking (using a changed threshold value), but deactivating rules 3i and 3c. Results for vel = 50, 40, 30 and 20 mV are shown in Fig 6 (for videos see Supplement). Although the starting configuration is apparently not well suited for these tests, stable patterns are reached after a few steps.

### Analysis of neural activity – The possible function of Intrinsic Oscillation

A large number of studies in insects and crustacea have shown that the central nervous system when completely deafferented and treated with pilocarpine, may show rhythmically oscillating activity of the motor neurons innervating flight muscles or leg muscles used for swimming or walking. These oscillations are generally assumed to result from central pattern generators forming the basis of a system controlling the rhythmic motor output observed in swimming, flying or walking animals. To challenge this hypothesis, in the following we use our hexapod simulator to test if alternative interpretations may be possible that show the same emerging intrinsic oscillations.

#### All legs deafferented

We will first deal with a thorough study concerning experiments with locusts (Knebel et al. 2017). In all these experiments, the three thoracic ganglia of a locust have been deafferented, as well as connectives to the suboesophageal ganglion and to the abdominal ganglia were cut (but see Knebel et al. 2018). With this preparation, a number of experiments have been performed by treating only selected ganglia with pilocarpine. The main results are that all six hemiganglia oscillate with a period of about 5 s and show in-phase coupling (recordings from depressor motor neurons), if all three thoracic ganglia were treated with pilocarpine. This result is supported by studies on stick insects *Carausius morosus* (Borgmann et al. 2009; Büschges et al. 1995). Corresponding data are also available for rock lobster *Jasus lalandii* (Clarac and Chasserat 1979 – recordings from retractor muscles, legs autotomized) and for crayfish *Pacifastacus leniusculus* (Sillar et al. 1987). In the study of Knebel et al. (2017) coupling between all hemiganglia remained in-phase if either only prothoracic ganglion, or only mesothoracic ganglion or both were treated with pilocarpine. When only the metathoracic ganglion was treated with pilocarpine, right and left hemiganglia of all three thoracic ganglia showed anti-phase coupling. Similar results have been reported for *Carausius morosus* (Mantziaris et al. 2017). Knebel et al. (2017) concluded that the oscillators of neighboring hemiganglia are all coupled via in-phase connections except the hemiganglia of the metathorax, which are coupled in anti-phase. Interestingly, the in-phase coupling observed corresponds quite well to coordination rule 5 (Dürr et al. 2004; Cruse 1985a) as introduced into Walknet (Schilling et al. 2007), see Fig 2, ocher units, 5i, 5c and Methods. In neuroWalknet the anti-phase coupling is realized as a connection between both metathoracic hemiganglia (rule 5ch, see Methods) coupled via inhibitory synapses (Fig 2, ocher units, upper left, lower left).

To what extent is it possible to simulate these results with our controller? As mentioned in Methods, we cut the velocity output to the motors thereby freezing all joint positions. Furthermore, the mode of each leg was set to stance and the inputs of the load sensors (marked by “pilo” in Fig 2) were set to high values, thereby simulating the effect of deafferentation and pilocarpine.

When, following the experiment of Knebel et al. (2017), in our simulation either only the prothoracic ganglion or only the mesothoracic ganglion was “treated with pilocarpine”, all six hemiganglia oscillated in-phase (Fig. 7B, C, activation of depressor motor neuron). The same result was found when all three ganglia were treated with pilocarpine (Fig 7A). When however only the metathoracic ganglion was treated, ipsilateral hemiganglia remain in-phase, but all contralateral hemiganglia oscillated in anti-phase (Fig 7D). These simulation results agree well with the findings of Knebel et al. (2017).

**Fig. 7 A-D.**
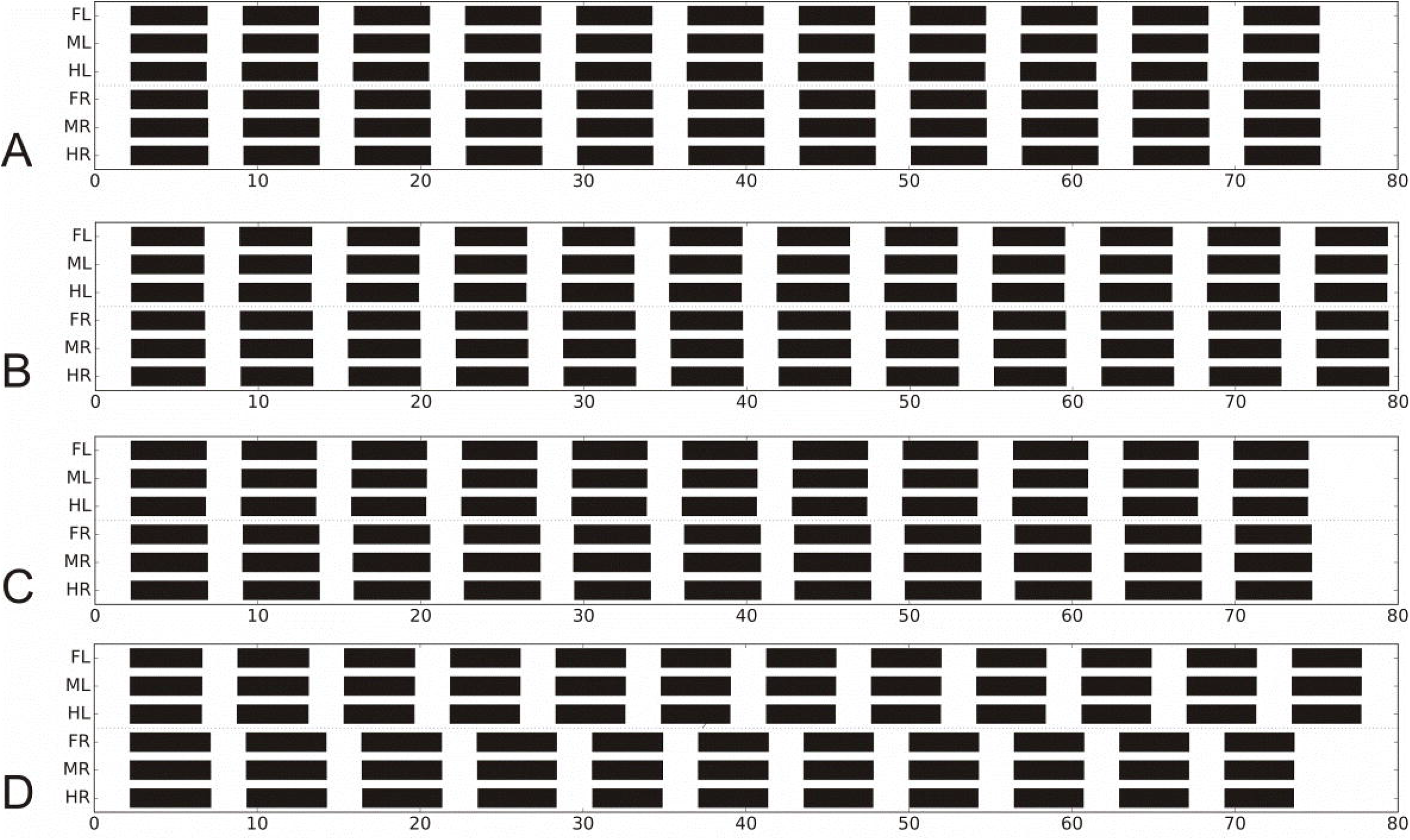
Simulation of experiments of deafferented locusts (Knebel et al 2017). Black bars show activation of depressor muscle output (> 0 mV). A) all thoracic ganglia treated with pilocarpine, B) only prothoracic ganglion treated, C) only mesothoracic ganglion treated, D) only metathoracic ganglion treated. Last five periods show stable state. Abscissa: time (s).

In the simulation given in Fig 7A, B, C, the antiphase influence occurring between the hind legs was apparently overridden by the stronger in-phase coupling of the other leg pairs. Note that influences from rules 1-3 were not effective in this situation as sensory input concerning leg position is fixed (and leg controllers were assumed to be in stance mode during the application of pilocarpine). Therefore, the critical effects resulted from rule 5 influences plus the extension assuming a contralateral inhibitory connection between the hind leg controllers (rule 5ch).

These simulations show that results of Knebel et al. (2017) can be reproduced although the leg controller does not contain any CPG that triggers the rhythmic movement during normal walking, i.e. controls patterns characterized as pentapod, tetrapod or tripod. Rather, due to the pilocarpine driven high excitation of both premotor units of the beta joint and sufficiently high mutual inhibition via the red units (PMN) connecting both branches (Fig 2), the four-unit premotor network started to oscillate with a period of about 5 s.

As rule 5 connections are applied to the alpha joint, too, independent oscillations were also observed in the retractor-protractor system, which was however not (yet) investigated by Knebel et al. (2017).

Coupling among both premotor units of the beta branches is assumed here in a way to produce levator and depressor activity with asymmetric ratio as observed in the animal (here we use a ratio of about 1: 2 duration). The connection weights could of course easily be changed to produce other, for example more symmetric relations.

#### Intact walking legs and deafferented legs

Interesting experiments have been performed by Borgmann et al. (2007, 2009) with *Carausius morosus*, where one intact leg, for example the right front leg, was walking on a treadmill while the other legs were amputated, i.e. deafferented only (Borgmann et al. 2007), or in addition treated with pilocarpine (Borgmann et al. 2009). The authors recorded movement and retractor activation of the walking leg (right front leg) and neuronal activity of the other ipsilateral legs. Recordings from the retractor motor neurons of middle legs and hind legs being deafferented and treated with pilocarpine showed that all ipsilateral hemiganglia were driven in-phase with the stance mode, i.e. retractor activation of the walking leg. Interestingly, corresponding results have been observed by Clarac and Chasserat (1979) studying rock lobster *Jasus lalandii* (legs autotomized).

How would neuroWalknet behave in this situation? Fig 8 shows the result when the left front leg (not the right one as used by Borgmann et al. 2009) was allowed to walk while all other legs were deafferented and “treated with pilocarpine” as used for simulation of the Knebel et al. (2017) experiments. In the simulation, all remaining ipsilateral hemiganglia oscillated in-phase and with the same period as the walking leg. This means that the oscillatory elements appear to be driven by the rhythm of the walking leg. As shown in Fig 8, the simulation further predicts that also contralateral legs should oscillate in-phase. To our knowledge such recordings have not been performed yet. Simulation of different walking velocities are provided in Suppl. Fig S5 indicating that the right hind leg appears to be less strongly coupled to the walking leg than all other legs. Note that this simulation may also be considered as an explanation of results of Wendler (1966), who observed stumps of amputated middle legs moving in-phase with the intact ipsilateral front leg walking on a treadwheel.

**Fig. 8.**
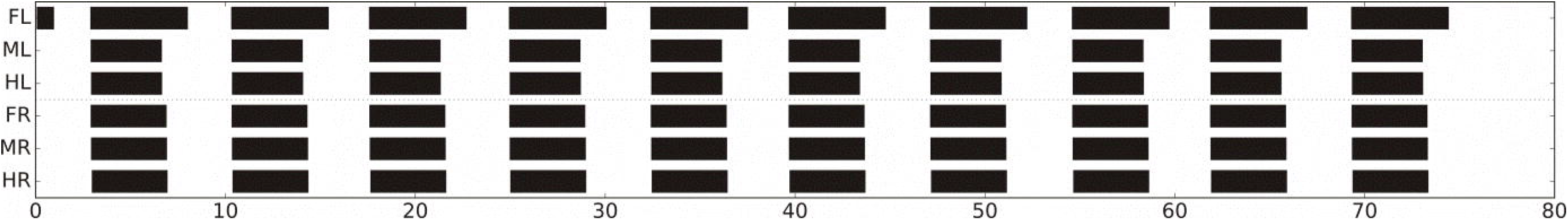
Simulation of experiments with stick insects, one leg (FL) walking on a treadmill (vel = 30 mV), while the other legs are deafferented and treated with pilocarpine (in Borgmann et al 2009, only ipsilateral legs were recorded). Black bars show retractor muscle output (> 0 mV). Velocity of walking leg 30 mV (for further velocities see Supplement Fig S5). Abscissa: time (s).

#### Standing legs in walking insect

Finally, a set of experiments will be discussed, where animals were supported upon a treadmill and only a few legs were able to walk, i.e. able to move the wheel. Different to the case of Borgmann et al. (2009), the other legs were not deafferented, but remained intact and were being placed to stand on a force transducer each being fixed aside the body. These standing legs often performed oscillating motor output (Cruse and Saxler 1980).

A preliminary experiment of this kind has been performed with the rock lobster (*Jasus lalandii*), but with only one leg on a force transducer (Cruse et al. 1983). The leg standing on the fixed force transducer showed rhythmic, backward directed forces in-phase with stance movement of the anterior neighboring walking leg. More detailed experiments were performed with stick insects *Carausius morosus*, where different configurations of walking and standing legs have been studied (Cruse 1980). It was however difficult to obtain clear results in some cases, probably because in these quite unnatural situations rules 1-3 influences and rule 5 influences are superimposed leading to irregular behaviors difficult to analyze. Further, simulation is difficult because the mechanical situation is different. Animals are supported above a treadwheel. In contrast, the robot walks with three or four legs, while the fixed legs are slipping on the ground which may lead to not well structured walking movements. We therefore focus on only those cases where at least three neighboring legs are walking. In these cases, a separation between rule 1-3 effects and rule 5 effects appears to be possible at least to some extent. Here we show results and simulation of three cases where the experimental results showed obvious maxima concerning appearance of posterior extreme positions (PEP) of walking legs and force maxima of standing legs.

First, we consider the case, where both front legs and both middle legs walk on the treadmill showing an about normal walking coordination (Fig 9A, simulation; Ai, biological data). Each of the standing hind legs showed strong forces pointing rearward in-phase with the retraction of the ipsilateral middle leg (note that motor output occurs earlier than leg movement). This result is recognizable in both behavior and simulation. How could this result be explained? In this situation, rule 3i-influences from middle legs to hind legs were not effective as the hind legs adopted a constant position in stance mode. Rule 5-influences from hind legs to walking middle legs had no effect when the middle leg was in swing mode. If the middle leg was in stance state, there was no specific change in leg velocity as all legs walking are mechanically coupled via the substrate. However, the standing hind legs appear to have received rule 5-influence from their neighboring middle legs elicited due to high friction in the middle legs walking on the treadmill. This effect was stabilized further by the anti-phase influence of rule 5ch active between both hind legs. Thus, there is an at least qualitative agreement between behavior and simulation.

**Fig. 9.**
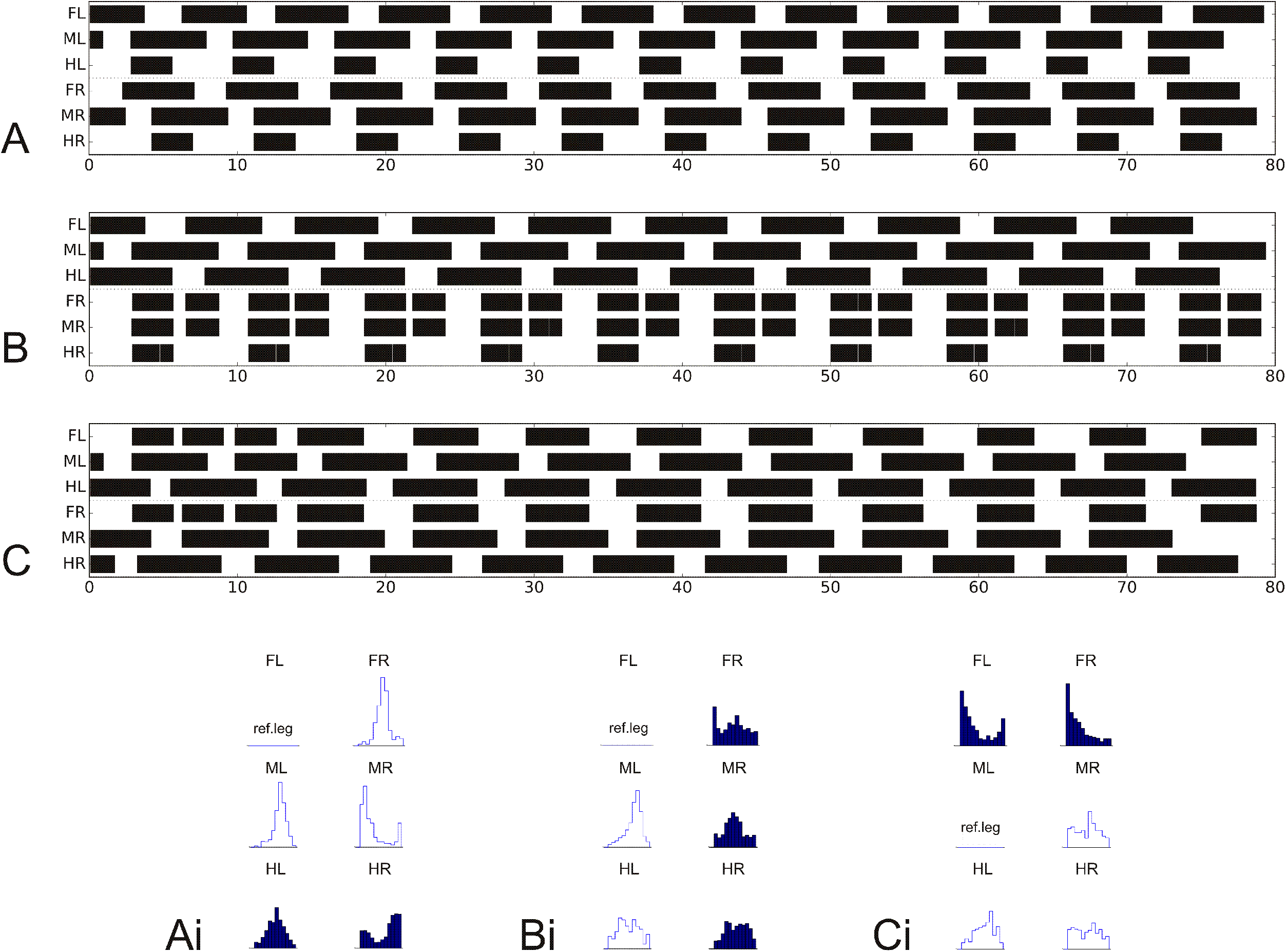
Simulation of experiments where stick insects walk tethered on a treadmill with selected legs standing on force transducer platforms. Shown are activations of retractor motor neurons (> 7 mV) over time (s) in A) to C). Corresponding experimental results (after Cruse and Saxler, 1980a and b and Schilling et al. 2007) are given in Ai, Bi, CI, respectively: Phase histograms of the beginning of the retraction in the walking legs (shown in white), force histograms of the maximum force in the standing legs (dark). Reference leg starting with the beginning of the retraction movement. In A) and Ai) both front legs and both middle legs are walking while both hind legs are standing. B) and Bi) shows left legs walking and right legs standing. C) and Ci) show both front legs standing, both middle legs and both hind legs walking (Cruse Saxler 1980, Schilling et al 2007). Walking velocity = 30 mV.

Another interesting case (Fig 9B) is given when the three left legs were walking on the wheel while each right leg was standing on its force transducer. The walking left legs showed a pattern somewhat slower than ideal tripod (behavioral data, see Fig Bi). The standing right front leg showed two force maxima, one in-phase with beginning of stance of the left, walking front leg, the second one being in-phase with force maximum of the right middle leg. The latter was in-phase with stance beginning of the left, walking middle leg (an in-phase influence to the right middle leg from the first maximum of the right, ipsilateral, front leg may exist but is not significant). Again, note that forces were somewhat earlier than leg positions.

How could this behavior be explained? The right front leg received rule 5-influence from the left front leg when starting stance which lead to its first (left) maximum. The second maximum was elicited by rule 5 input from the ipsilateral, standing middle leg, which in turn resulted from the beginning of stance of the left, walking middle leg. An ipsilateral rule 5-influence from the right front leg may exist, but if so, might have been suppressed by less focused rule 5 influences from the standing (right) hind leg. The bimodal force output of front leg and middle leg is clearly visible in the simulation. The unclear force distribution in the right hind leg may result from opposing influences, first the ipsilateral rule 5 influence from the right middle leg and second the weaker anti-phase influence via rule 5ch from the left hind leg. There is a broad maximum in the standing hind leg (Fig 9 Bi), but it is not clear if this maximum agrees with the one observed in the simulation.

The interpretation of the case shown in Fig 9C, Ci – both front legs are standing on a force transducer – seems more difficult, as there is only weak coupling between right and left walking legs. This may result from lacking of 3i-influences from front legs to middle legs which includes lacking 2c and 3c coupling via the front legs. A phase shift between contralateral neighboring walking (middle and hind) legs is also observed in the simulation. In contrast, both standing front legs showed strong in-phase coupling, being recognized in behavior and simulation (Fig 9C, Ci). Extreme in-phase coupling between front legs had been reported by Cruse and Saxler (1980, their Fig 3) suggesting a quite strong rule 5-influence, at least in a situation where both front legs try to pull the body forward. The asymmetric distribution of the force maxima in the front legs results from the fact that left middle leg was used as reference in this evaluation.

Taken together, the behavior observed in quite unnatural situations of standing legs in a walking animal appears to be, at least in principle, in accordance with the properties implemented in neuroWalknet.

## Discussion

How to control natural behavior that requires more than just controlling simple reflexes is still a question under debate. ‘Natural behavior’ means that the agent has to deal with a complex, unpredictable environment as well as with a motor system that is characterized by a high number of degrees of freedom (e.g. joints). These DoFs may be arranged serially or in parallel and usually include redundant DoFs, i.e., an arrangement where several solutions are possible to reach the same behavioral goal. Hexapod walking and climbing on unstructured substrate represents such a case. Here, we propose an artificial neural network, neuroWalknet, a decentralized controller which is suited to approach such problems and which represents a testable hypothesis as to how walking behavior as studied in insects may be controlled. The neuronal basis of the neuroWalknet approach complements the original Walknet approach with a detailed neuronal realization based on an antagonistic structure.

This neural-based control approach was tested in different behavioral settings. As has been shown neuroWalknet is able to produce footfall patterns that include “tripod” patterns, “tetrapod” patterns, “pentapod” patterns as well as various stable intermediate patterns. These patterns form a continuous multitude as has been observed in stick insects (Graham 1972) and in Drosophila (Wosnitza et al. 2013; DeAngelis et al. 2019) and to some extent in cockroaches (e.g. Hughes 1952; Bender et al. 2011). Any disturbances provided by the environment add to the variability of patterns. However, if no such disturbances are present, a stable pattern will be reached that depends on the value of the global velocity chosen. Eventually, mirror image patterns are possible with phase coupling among contralateral legs deviating from phase values of 0.5. Patterns recently described as non-canonical by DeAngelis et al. (2019) can be observed, too (e.g. Fig 3C). Furthermore, neuroWalknet allows for backward walking and negotiation of curves. A leg controller of this type may be classified as a “free-gait” controller (Schilling et al. 2013a) because the network does not rely on motor memories representing explicitly defined specific patterns. Rather, the behavioral patterns observed reveal emergent properties of a decentralized neuronal architecture, which easily allows for starting or interrupting a walk or changing the velocity. Overall, neuroWalknet shows the same behavioral variability and adaptivity as the original and well-established Walknet approach.

In addition, the controller allows to simulate results of neurophysiological studies generally assumed to support the highly influential concept that CPGs are required to control the rhythmic movements of walking legs. We show that a number of experiments can be explained without relying on this concept.

In general, neuroWalknet is a dynamical systems approach from which activities emerge in interaction between different decentralized components of the control system and with the environment mediated through the body and its properties. This can be observed on many different levels: while apart from very fast walking (beyond 5-7 steps/s, see below and Supplement) in the intact animals no intrinsic oscillations are observed on a neuronal level, gaits are the resulting behavior. neuroWalknet provides an approach in which these different patterns emerge from interaction on these different levels. In this section, we will focus on these levels subsequently and start from bottom up:

- first, starting with the neuronal level and addressing how neuroWalknet is in agreement with findings on the neuronal structure of such control system in insects and where it deviates,
- second, dealing with control on the level of the individual leg, and,
- third, addressing coupling between legs and how emerging coordination can be explained through the system including temporal patterns on the neuronal level from deafferented animals.

### A) Neuronal Counterparts

neuroWalknet may provide a functional counterpart for some neurons described in neurophysiological studies, thereby representing a hypothesis as to how these neurons might be embedded into a complete system.

#### Representation of Walking Parameters on the neuronal level

The four motivation units required to decide between forward mode and backward mode and to stabilize the latter (dark gray units) may correspond to neurons described by Bidaye et al. (2014) and recently successfully tested by DeAngelis et al. (2019) in *Drosophila*. These authors found two ‘moonwalker descending neurons’ (MDN) on either side in the protocerebrum, which may function in a way corresponding to the rightmost dark gray unit (“backward”) activated by antennal stimulation (Fig 2), and may stabilize backward walking, while the ‘moonwalker ascending neuron’ (MAN) may correspond to the upper central dark gray unit, which stabilizes backward walking by inhibiting the leftmost dark gray unit (“forward”) responsible for forward walking.

Another neuron (Fig 2, leftmost white unit) is crucial for the controller, as it represents the current walking velocity. This unit may functionally correspond to a group of neurons observed in crickets (Kai and Okada, 2013) that fire closely related to walking velocity. Further candidates have been found in the central complex of cockroaches (Bender et al. 2010).

#### Neural representation of Searching Movements as part of an Adaptive Swing Movement

In stick insects, searching movements of a leg are observed if the leg at the end of the swing movement is not stopped by ground contact (Dürr 2001). Frequencies of searching movements are about 3 Hz (and about 10 Hz in *Drosophila*, Berendes et al. 2016). In both cases, these values correspond to that of the uppermost leg frequency during fast walking. Starting from our initial assumption (see Introduction), one could ask if such searching movements represent a distinct behavior which would be addressed separately on the level of motor selection, or if searching movements just simply are part of an adaptive swing behavior. In an excellent study concerning the control of searching movements, Berg et al. (2015) investigated the contribution of non-spiking inhibitory interneurons during stance and during swing as well as during performance of searching movements in a restrained middle leg and found two antagonistically active groups. One group is hyperpolarized during flexor activity, while neurons of the other group are depolarized during flexor activity. Two neurons, I1 and I4, each belonging to one of either group, have been studied in more detail. I1 belongs to the former group and is active during stance, I4 belongs to the latter one and is active during swing movement and during searching movements in *Carausius morosus*. Berg et al. (2015) could show that swing and search is stopped either by foot contact or by hyperpolarization of I4. Artificial neurons of this type are found in neuroWalknet, too, whereby the lower light grey unit in Fig 2 may correspond to a neuron like I4 and the upper light grey unit may correspond to a neuron like I1 (no information is given as to the effect of neuron I1). If this interpretation is correct, an additional neuronal unit representing a searching movement would not be necessary, contrary to what has been postulated by Berg et al. (2015) and tested in a simulation by Szczecinski et al. (2017). Therefore, swing and search behavior may result as an emergent property from the network already required for basic walking. In this view, the oscillations observed during searching result from the properties of negative feedback loops controlling the joints during swing as given in neuroWalknet (not simulated) and as has already been proposed in a simulation by Dürr (2001).

If at the end of swing, during touch down, the tarsus does not find efficient adhesion to the substrate, the leg may react by lifting the leg up and down similar to that of a searching movement. This behavior corresponds to what has been described as “short steps” (Theunissen and Dürr 2013; Bläsing and Cruse 2004) and might be termed short step A (for anterior) in contrast to short step P (for posterior, see Methods and Supplement). As short step A may be caused by the same structure as used for searching, but under specific environmental conditions, we did not simulate this behavior, in contrast to short step P (see Supplement).

#### Ring network and Central Complex

A hypothetical ring net as proposed here to exist in each thoracic hemiganglion (Methods, Fig 10) shows parallels to the ring network forming the central complex, a prominent structure in the insect brain. As interpreted by Stone et al. (2017) and others, this structure may serve for vector navigation. To this end, it receives input used to record the direction of current body axis in a geocentric coordinate system and information concerning the currently desired walking direction theta measured in a body-fixed coordinate system. These signals are projected to the noduli. Stone et al. (2017) postulate that the noduli provide a vector representing the walking direction which is given to the motor system (for neurophysiological data see also Martin et al. 2015). Our proposed ring net (Methods, Fig 10) receives as input angle theta and the current leg position represented by angle alpha, and provides as output a direct influence on alpha angle and on gamma angle of the corresponding leg, as described in Methods, representing the components forming the leg trajectory. This means, when compared with the central complex, the leg ring network might represent a similar, but much simpler structure where layers 2 and 3 correspond to the input to the protocerebral bridge and layer 4 and 5 plus units O1 – O4 (Fig 10) represent the functional aspect of the noduli.

**Fig. 10.**
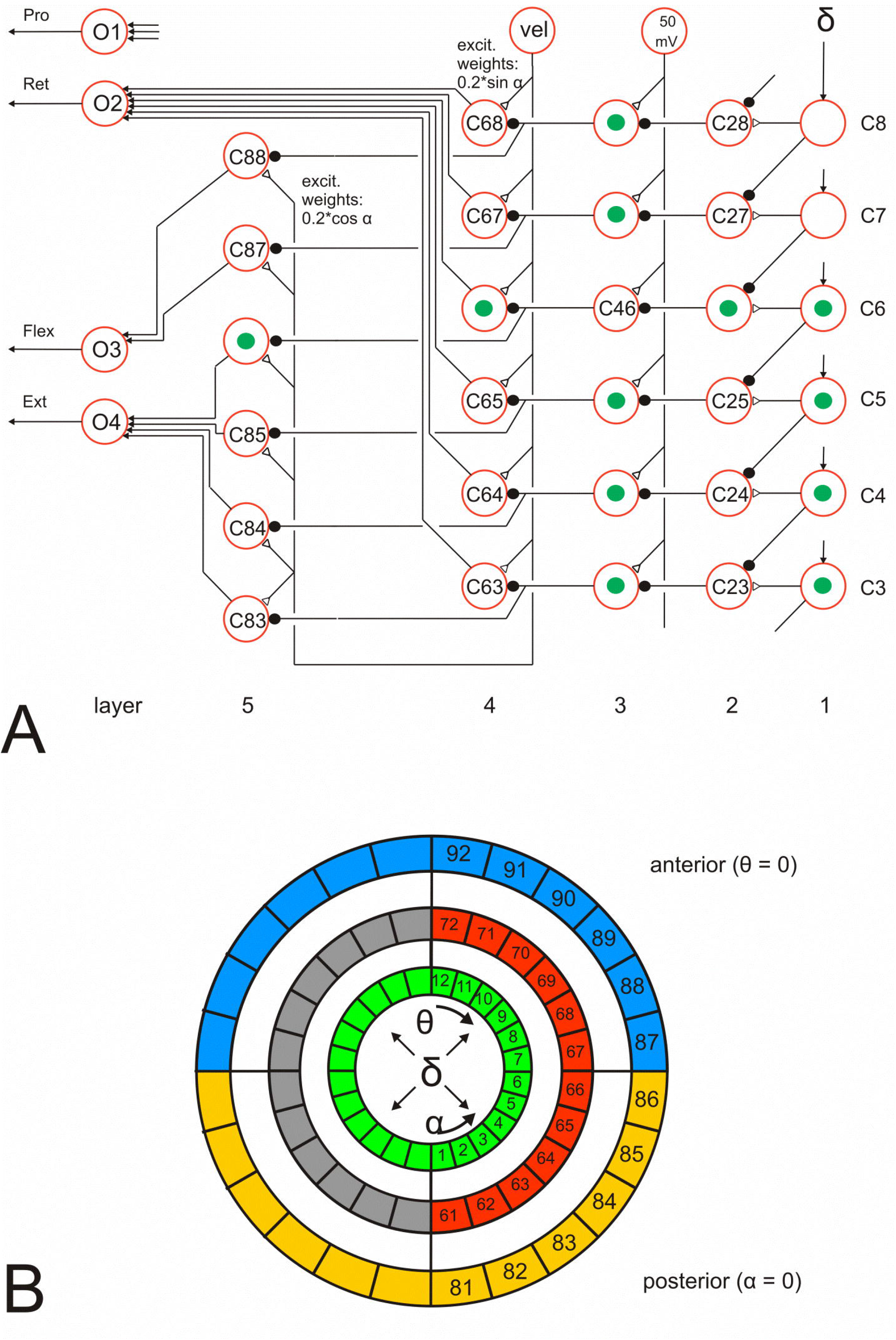
Ring net. A) Angle delta (leg position, angle alpha, and walking direction, theta) determines the contribution of alpha joint and gamma joint during stance. Column 1 (right) represents angle delta in spatial coding. Depending on desired velocity of the leg (vel) and angle delta the motor output for the retractor (O2), for flexor (O3) and for extensor (O4) are computed. Protraction (O1) is not yet implemented. Units marked with green dots show, as an example case, activation of the net by a delta value input of spatial code 6 (e.g. theta = 0 degrees and alpha = 25 mV).). B) Organisation of ring net, schematically. Input: layers 1 – 3, green. Input delta = alpha – theta. The analog value delta (in mV) is transformed to the spatial coding version (Fig 2, box “spatial coding”), which is then given to the circle marked green. Output: Layer 4 (red, grey) controls the alpha joint (red: retractor, unit O2, green: protractor, unit O1, not used in the current version). Layer 5 (blue, yellow) controls the gamma joint (blue: flexor, unit O3, yellow: extensor, unit O4). Letter C (for numbers of circle units, Fig 10A) is not shown in Fig 10B.

One may therefore speculate that the ring net, assumed to be realized in each thoracic hemiganglion, may be an evolutionary early, simple network suited for controlling the individual legs. The central complex may have arisen later as a merged version of right and left ring networks thereby forming a centralized and more sophisticated structure required for controlling the walking direction of the animal to be used for vector navigation (Stone et al. 2017, Greene et al. 2017). Information concerning desired walking direction provided by the central complex may then be given to each of the six ring nets.

### B) Individual leg control

As one key novel property, neuroWalknet is based on detailed neural control networks and includes antagonistic structures. This allows to analyze and compare different hypotheses on the neuronal level.

#### Control of the movement of an individual leg

For their model, Daun-Gruhn and Büschges (2011, their Fig. 3) introduced an interesting distinction between “timing” and “magnitude”. Timing is controlled by a WTA structure, in their case a CPG that depends on internal commands (e.g. global velocity) and on sensory signals. Magnitude concerns the determination of the strength of motor output depending on sensory feedback. In principle, this concept is also applicable to the structure of neuroWalknet. The main difference between their approach (see also Rubeo et al., 2017) and the approach presented here is that in neuroWalknet there is only one controller for timing, represented by the four light grey units, that concerns all three joints of a leg, whereas in the model of Daun-Gruhn and Büschges (2011) and of Rubeo et al. (2018) each joint has its own “timer”, the CPG of the respective joint. In neuroWalknet, Fig 2, the leftmost (“swing”) and the rightmost (“stance”) light gray units receive input from sensors (leg position, load) of the own leg and, via coordination signals, from neighboring legs, which in turn depend on the positions and on the state of units “swing” and “stance” of those legs as well as on the global velocity signal. In other words, these (light grey) WTA nets, one for each leg as used in neuroWalknet, rely on various sensory signals and the global velocity signal.

The other two units of this winner-take-all (WTA) net (upper, lower center) provide negative feedback required for a WTA structure, the parameters of which are chosen to prohibit oscillations, that may otherwise lead to instable walks. Mutual inhibition between units “swing” and “stance” is indeed necessary for two reasons: When the hexapod starts with an uncomfortable leg configuration, both units might receive contradictory signals which requires a decision as to which of both units should be activated. But the inhibitory connections are also necessary during stable walking, because during walking there are time periods where none of both units receives input. In this case, the mutual inhibition system acts as a short term memory (or as attractor states, see Conclusions) maintaining the actual state until new input is arriving. This, in turn, means that a feedforward inhibition may not be sufficient. Rather, a recurrent inhibition is required.

Rubeo et al. (2018) use CPGs for each joint as a basis to control walking rhythms, comparable to the approach by Daun-Gruhn and Büschges (2011). They connect neighboring legs by direct coupling of CTr flexor muscles MNs (corresponding to the levator muscle of stick insects) through coordination rules 1 - 3. This is an excellent study, which is very much related to our approach, but is focusing on simulation of fast walking cockroaches. Compared to Hector, the authors applied a much more sophisticated simulation of, in this case Hill-type, muscles. But as they deal with cockroaches, their focus is solely on tripod pattern with a given velocity (5 steps/s). Therefore, their approach appears complementary to ours as we focus on slow walking for control of the single leg, and instead apply the WTA net (marked by light grey units) to control leg movement. As a consequence, in neuroWalknet control of muscles is not strictly connected to the higher-level control represented by the light grey units. Therefore, different controller types can easily be used for either swing or stance. But also within stance state or swing state variable application of antagonistic muscles is possible (e.g. Fig S 2A, e.g. alpha joint, gamma joint). As a consequence, implementation of avoidance reflexes or short steps as well as searching behavior is easy within neuroWalknet, because a local change of antagonistic muscles does not influence the overall state of the leg. Furthermore, as no CPGs are applied on this level, our structure strongly simplifies height control, negotiation of curves and dealing with disturbances, as no dynamical oscillatory systems have to be constrained. This is particularly obvious for starting and stopping. As shown by Tóth and Daun (2019), in their system based on coupled oscillators various conditions have to be fulfilled to overcome the dynamic behavior of such a system (related problems are described by Rubeo et al (2018). This is in strong contrast to our solution, where simple setting of global velocity from zero to a positive value or back is sufficient. This means that application of the WTA structure increases the adaptivity of the behavior dramatically and shows that for walking – apart from running – CPGs are not only not necessary, but may cause negative effects. Different approaches on legged robots apply simple CPG approaches for temporal coordination [Kalakrishnan et al., 2010; Ijspeert, 2014]. While these excel at spatial coordination in rough terrain, recently Bellicoso et al., [2018] argued that—for their case of a quadruped robot—climbing and changing environmental situations also require adaptation on the temporal level which appeared difficult when dealing with fixed gaits.

#### Levator Reflex

The levator reflex in *Carausius morosus* is elicited if during swing movement the leg touches an obstacle. In this case, the leg shows a brief retraction accompanied by a levation, followed by protraction to continue the swing movement (e.g. Schumm and Cruse 2006, Ebeling and Dürr 2006). This reflex can easily be simulated if a tactile stimulus to femur or tibia is represented by a fictive sensor and the stimulus is given via a neuron to the retractor branch and to the levator branch as has been shown by Schumm and Cruse (2006). Such a levator reflex is implemented but no results are shown.

### C) Interleg Coordination

In the following, we will further discuss specific aspects concerning interleg coordination in more detail.

#### No Central Rhythms required for slow walking

As argued above, gaits are emerging in neuroWalknet, which represents a free gait controller. This assumption is often contrasted with the notion of fixed rhythmic patterns that are assumed to be created by Central Pattern Generators. neuroWalknet can also produce central rhythms as observed in a number of neurophysiological studies on insects (Knebel et al. 2017; Büschges et al. 1995; Mantziaris et al. 2017) and crustaceans (*Jasus lalandii* – Clarac and Chasserat 1979; crayfish – Sillar et al. 1987), but without relying on dedicated Central Pattern Generators that are used to control normal walking. The simulations show that in the case of neuroWalknet these rhythmic oscillations can be explained by a combination of known coordination rules (rule 5, Dürr et al. 2004) including an expansion proposed by Knebel et al. (2017) and the effect of deafferentation and pilocarpine being applied in these experiments. Furthermore, several experiments can, on a qualitative level, be explained in which either neurophysiological recordings in deafferented legs are combined with legs walking on a treadmill (Borgmann et al. 2009) or where intact legs standing on a fixed force transducer are combined with legs walking on a treadmill (Cruse 1980). In the latter case, oscillatory motor output can be observed although no pilocarpine has been applied. Interestingly, in principle corresponding results have also been observed in crustaceans (Clarac and Chasserat 1979; Cruse et al. 1983), arguing for a general relevance of a controller structure as proposed here.

These results provide an alternative explanation for quite a number of experimental findings which challenges the generally accepted view that CPGs observed in different insects or crustaceans do represent the basis for the control of multi-legged walking.

#### Testing other types of coordination between legs

Concerning coordination influences between legs, Tóth and Daun (2019) postulate that various direct coupling connections between hind legs and front legs are necessary to establish coordinated locomotor pattern. There are, however, no experimental hints supporting such an assumption. Their conclusion appears to be valid only under the assumption that CPGs are indeed responsible for control of walking. neuroWalknet does neither contain direct coupling between front legs and hind legs, nor does it use CPGs to control walking rhythm, but nonetheless allows the control of different patterns plus stable intermediate patterns as observed in stick insects and *Drosophila*.

#### Interleg coupling dependent on walking velocity

Along a similar line, Borgmann and Büschges (2015) report that coupling is weaker in slow walking insects compared to fast walking. Indeed, the simulation with neuroWalknet shows that for very slow velocities (e.g. velocities of 10 mV or less) and arbitrary starting configurations it may take more time until a stable pattern is reached. However, as above, weak coupling observed on the behavioral level does not mean that neuronal parameters, i.e. synaptic weights, are changing depending on velocity. In neuroWalknet this effect is simply caused by signals that – sent via the coordination channels – occur more rarely in slow walking. Therefore, more time is required until a stable pattern is reached after a given disturbance.

DeAngelis et al. (2019) propose a very simple network for interleg coordination in *Drosophila* with contralateral legs using only phase value of 0.5, ipsilateral legs using only forward directed influences and all parameters are not dependent on walking velocity. For stick insect (*Carausius morosus*) detailed studies show that ipsilateral coupling operates in both directions and contralateral legs show phase values that differ from 0.5 during tetrapod walking. These data and those from Graham (1972) further show that both ipsilateral coupling and contralateral coupling depends on velocity. Therefore, the coordination influences in *Drosophila* may be simpler than those found in *Carausius*, or studies that reveal more detailed behavior as performed with *Carausius* (review Cruse 1990, Schilling and Cruse 2013a) are lacking in *Drosophila*.

Interleg coupling dependent on walking direction: As illustrated in Fig 6, walking backward shows stable footfall patterns similar to those observed during forward walking. This seems to be in contrast to the results of Pfeffer et al. (2016), who stated that coordination in backward walking ants is weaker than in forward walking. The reason for this weak coordination may be that in these experiments ants had to carry higher load when walking backwards which provides irregular sensory input that may disturb the controller or, that the ants simply adopt a lower velocity (see above). So, actual coordination observed on the behavioral level is weaker indeed, but this does not necessarily mean that neuronal parameters responsible for interleg coordination have changed.

#### Backward walking

Szczecinski et al. (2018) could show that a more statically stable version of hexapod backward walking can be observed when the direction of the ipsilateral sequence “rear-middle-front” of legs swinging is reversed to “front-middle-rear”. This sequence has indeed been observed in *Drosophila* (Bidaye et al. 2014) and in *Aretaon asperrimus* (Jeck and Cruse 2007), but not in *Carausius morosus* (Graham and Epstein 1985). Here we adopted the simple version that leads to the *Carausius* pattern. The alternative pattern, which may meet the situation of other insects, would at least require introduction of additional, inverted versions of rules 1 and 2i, that operate from any leg to its posterior neighboring leg. Qualitative behavioral observations indicate that *Carausius morosus* appears to avoid spontaneous backward walking, which suggests that this species may not be equipped with further specific rules allowing for more “comfortable” backward walking, in contrast to species like *Aretaon* or *Drosophila*.

#### Curve Walking

One well documented case of curve walking (Dürr and Ebeling 2005) allowed us to test the ability of the controller to negotiate curves (Supplement Fig S 3). We have introduced a new neuronal element, “local(theta)” (Fig 2, box local(theta)), that, together with the ring network, is able to allow for negotiation of curves. However, this was only done for that specific turning angle theta, because only for this case sufficient data were available. In principle, it would be easy to implement a general function that, depending on theta, provides data for swing (AEP) and stance (local leg velocity, local turning angles). Such a function should however be based on biological data which are currently not available yet.

#### Stability of behavioral patterns

Studying the stability of behavioral patterns, as in our case represented by the footfall patterns, is difficult as the system may be characterized by showing bifurcations where small disturbances may lead to large effects. Nonetheless, apart from choosing specific starting configurations we have performed systematic tests with different legs, different velocities and different durations of disturbances. Detailed results are, however, not shown here, as only general qualitative observations are possible (but see specific examples in Figs 3, 4, 6). In all cases studied the disturbance only affects a couple of steps after which a stable pattern was reached again. Further, in general, as to be expected, lower walking velocity leads to smaller deviations and small disturbances lead to comparatively small deviations from the stable pattern. Overall, these observations are in full agreement with the many observations and different analyses of patterns found in the former Walknet approach (Schilling et al. 2013a for a review).

#### Very fast walking

Cockroaches, one of the most intensively studied insect as concerns walking, are often characterized as typical tripod (or “double tripod”) runners. However, as already reported by Hughes (1952) and Delcomyn (1971) they also use non-tripod gaits. This is supported by results of Bender et al. (2011), Full and Tu (1990, 1991), Ting et al. (1994) who found, in cockroaches, a continuum of walking velocities from zero to 0.5 m/s, corresponding to a frequency of nearly 15 steps/s. The latter means that the duration of a swing movement or a stance movement might be as short as 30 ms. Therefore, due to sensory delay unavoidable in neuronal systems, sensory feedback cannot be used beyond a given step frequency (for insects like cockroaches *Periplaneta americana* or *Blaberus discoidalis* of about 7 steps/s, Sponberg and Full 2008; Delcomyn 1991; Zill and Moran 1981). This suggests that at least beyond this limit a controller may be installed that has to operate without sensory feedback, i.e. requires intrinsic (e.g. neural) pattern generators to control rhythmic movement. The generally proposed solution to this problem is to apply CPGs, which may possibly indicate a further evolutionary development allowing for very fast running.

In the Supplement we show how a corresponding expansion can indeed be implemented in neuroWalknet (Fig. S7). Importantly, this expansion exploits (i) already existing structures to introduce CPGs for controlling rhythmic movement, (ii) an intraleg coupling inspired by results of Büschges et al. 1995, and (iii) an interleg coupling based on results of Pearson and Iles (1973). As for all open-loop control approaches this requires a form of stabilization to counter drift. In CPG-based approaches this is often realized through introducing mechanical limits (which may include nonlinear properties of muscles) on joints or some modulating sensory feedback (Kukillaya and Holmes 2009). This allows for stable walking controlled by cyclic motor output in our case as well. High-speed bipedal running observed in cockroaches (Full and Tu 1991) may represent an even later development, which is however not considered here.

To summarize, many behavioral studies and simulation models take central rhythms as a basis of walking and see CPGs playing a causal role. But this concept is based on mostly neurophysiological studies and on an only very limited number of behavioral studies under usually quite restricted conditions. For more natural behaviors that require a much higher adaptivity and are characterized by a much higher variability, such approaches seem ill-suited as they assume that the role of sensory information gathered during locomotion only plays a subordinate role. Indeed, such a secondary role of sensory signals required to overcome sensory delay appears an important feature for running in contrast to walking. Such a distinction of heterogeneous contributions of sensory information on motor control depending on behavioral state is assumed as well in mammals. In a recent study, Clancy et al. (2019) found differing connection strength from sensory areas to motor control areas depending on behavioral state and related to locomotion leading the authors to the conclusion that locomotion might shift cortical control networks more and more into a regime giving greater relevance to sensory feedback for slower velocities. Adaptive locomotion as found in slow walking, negotiating curves and climbing through twigs seems to require a lot more sensory information for stable behavior. Therefore, neuroWalknet as proposed here agrees well with these findings.

## Conclusion

neuroWalknet can reproduce a wide variety of behaviors which is unsurprising as main organizing principles are taken from the original Walknet approach. However, the fine grained structure of neuroWalknet using artificial neurons as basic elements allows in addition to simulate a number of experimental data concerning specific neurophysiological studies, never reached by a former approach.

Importantly, the overall behavior emerges from the interaction between various decentralized ‘motor memory’ elements and the environment mediated through the body. These motor elements can be seen as organized on different levels. Basically, there are six memory elements, one for each leg. Each of them contains three smaller motor memories, one for each joint. The latter in turn can be subdivided into elements contributing to swing mode or stance mode. On a higher level, the controller can switch between states as are “forward walking” and “backward walking”. Another internal state is given by the “Run” mode (Supplement). Even higher levels could be implemented as are, for example, states like “Searching for food” or “Homing”. The underlying neuronal architecture, called ‘motivation unit based columnar architecture’ (for detailed explanation see Schilling et al. 2013b, Cruse and Schilling 2015) can further be extended to represent symbolic elements, being combined with (sensory) motor elements. Symbolic elements may not be relevant for a neural structure of stick insects, but there are strong arguments that such elements may be required in other insects, for example honey bees or bumble bees (Loukola et al. 2017; Menzel et al. 2007).

These different memory elements are controlled and stabilized by a recurrent neural network (RNN) termed motivation unit net that allows for various stable attractor states, which can control quite complex behaviors. To this end, these memory elements are bound together in variable combinations by the motivation unit network and by the network representing the coordination rules.

Simulation of a broad range of experimental behavioral and neurophysiological data is possible as neuroWalknet adopts a view different from generally proposed assumptions because the contribution of CPGs is interpreted in a fundamentally different way. Following our interpretation the basic controller concerns slow walking, with strong impact on walking on irregular substrate including the requirement of developing higher forces in longitudinal direction. These properties are realized in both insects and crustaceans, whereas CPG-based running is assumed to having been developed only later. We believe that this view might stimulate comparative studies going beyond the traditional concentration on a small number of species only.

## Materials and Methods

The simulator consists of two parts, one for the neuronal controller and one for the body of the hexapod robot Hector, which exists as a hardware version and as a dynamic simulation (implementations are publicly available: dynamical simulation environment is realized in C++ and based on the Open Dynamics Engine library, for code base of simulator see https://github.com/malteschilling/hector; the neuroWalknet controller has been implemented in python (version 3), for python code base see https://github.com/hcruse/neuro_walknet). Here we use the dynamic simulation. In the following, we will, first, describe the simulation environment, and second, the realization of the neural controller including the neuron model. Finally, we will give details about how specific experiments were realized using the simulation.

### The simulated robot Hector

The robot Hector contains six legs with three active joints each and is designed to reflect important aspects of the geometry of stick insects with a scaling factor of 1:20 (Fig 1). The three joints are called alpha joint for the Thorax-Coxa joint, beta joint for the Coxa-Trochantero-femur joint, and gamma joint for the Femur-Tibia joint. Each joint is constructed as a hinge joint (which is a simplification, because in stick insects the basic joint has more than one degree of freedom, and in other insects additional joints can be found). The joint rotation axes are oriented as is the case in stick insect *Carausius morosus*, which means that the axes are not orthogonal, but show different inclinations for each leg (Schneider at al., 2014). Mechanoreceptors recording joint position and higher derivatives exist in each joint. In insect legs hair plates or hair rows are found in the Thorax-Coxa joint and the Coxa-Trochanter joint. The prominent sensor in the gamma joint is the femoral Chordotonal organ. However, in all joints there are various other sensors, but it is not known in detail in which way they contribute to the motor control system. Therefore, as a further simplification, we assume that, for each joint, there are antagonistically structured position sensors (Schumm and Cruse 2006; Szczecinski et al. 2013; Cruse 1980). This means that each joint is assumed to contain two sensor structures each of which monitors the full range of joint position, but with increasing excitation in opposite directions, i.e., we deal with redundant structures.

For simplicity, the three leg segments (Coxa, Trochanterofemur, Tibia) are identical for all legs. This differs from the insect where in front legs and in hind legs the Tibia (and to a smaller extent the Femur) is longer than that of the middle leg. For technical reasons the joints have a smaller range of movement compared to that of the animals. Therefore the maximum size of an average step used here is about 33 cm compared to the 40 mm in stick insects, defined by average values of anterior extreme position (AEP) and posterior extreme position (PEP) as used during normal, undisturbed walking. Different to the animals, there is no tarsus, and therefore there are no adhesive structures at the end of the foot. Another difference concerns the position of the center of mass (COM), which in the stick insect is placed between the Coxae of both hind legs, whereas in Hector it is placed slightly in front of the Coxae of both middle legs (Paskarbeit, 2017).

#### Measuring the Position of the Joint and Estimating the Torque

The simulator and the robot are equipped with joint position sensors that are used for a negative feedback position controller. Neither the robot nor the simulator does (yet) contain explicit force sensors. However, loading or unloading a leg may provide important information. As there are no explicit load sensors given in robot Hector, we recorded the desired position of each joint and the actual position reached after the elastic element of the motor. The difference is used as an approximation of the torque produced by the motor (see Fig S 2B).

#### Joint Actuation

According to results of Mu and Ritzmann (2005), Tryba and Ritzmann (2000a) and Watson and Ritzmann (1998) in cockroaches (*Blaberus discoidalis*) the motor neurons control joint velocity, which is the case for the motor neurons used here, too (Fig 2, MN). Examples are given in the Supplement (Fig S2A). The controller of the motor receives these velocity signals as set points for a PID controller. PID control for moving velocity of leg joints is supported by Levy and Cruse (2008), who found that in a standing stick insect the torques of the leg joints may have changed significantly although position of joints is not changed at all, i.e. velocity is set to zero.

The torques developed to reach the desired velocity can approximately be represented by the deviation between the desired joint position and the real joint position which differ due to an elastic element implemented in the motor (Schneider et al. 2014). In the simulation, the spring constants for the beta joint (see below) vary following the relation 8.5:4.5:8.5 for front leg, middle leg and hind leg, respectively. Middle legs are equipped with a softer spring to minimize pitch movements (i.e. tipping to front or back) when the middle legs are in stance mode.

The robot simulator, as the physical robot, is run by an update rate of 100 Hz, i.e., an update time of 10 ms. The motor output is adjusted in such a way that the behavior (duration of swing and stance) shows a ratio of about 10:1 compared to that of stick insects.

### The neural Controller

We will explain the neural controller starting with the employed neuron model, which is one major change compared to Walknet. The leg level and the individual joint control will be explained second. Third, coordination between legs will be addressed.

#### A) Neuron Model

The neural controller consists of artificial neurons. The dynamics of these are given by a simple differential equation as proposed by Hodgkin and Huxley (1952). We follow here the formulation of Szczecinski et al. (2017):

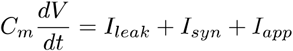

For the discrete time domain this is realized in the python implementation as

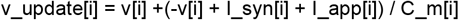

where [i] denotes the index of the neuron, v the voltage at time t-1, v_update the updated voltage at time t, I_syn the input from synaptic connections. I_app external (e.g. sensory) input (I_leak is set to zero normally, but see below). This means the dynamics show low pass filter properties with a time (membrane) constant C_m[i] of 4.5 ms, if not stated otherwise (for details on the implementation see code base). Update time is 1 ms (= 1000 Hz). Different to the classical Hodgkin-Huxley equations, the neuronal units currently used show a piece-wise linear characteristic with a lower threshold of 0 mV and an upper bound of 50 mV. Synaptic inputs are summed up linearly and, separately for excitatory input and inhibitory input, are clipped at 80 mV. As a further simplification, I_leak was set to zero. Only selected units are in addition equipped with a leak to allow for adaptation to a constant input, i.e. these units show properties of bandpass filters (marked HPF in Fig 2). To approximate the neural dynamics, high time constants (e.g. leak of 0.0001) are used when voltage approaches the resting potential (0 mV), whereas small time constants (leak = 0.1) are applied to recover from values below 0 mV. Following Dale’s law (Eccles et al. 1954), any unit is either inhibitory or excitatory, i.e. a unit is equipped with either excitatory output synapses or inhibitory output synapses.

#### B) Leg Control

Each leg is controlled by a network of about 164 units (plus ring net, Fig 10) and has an identical architecture. A leg controller consists of three subnets (Fig. 2, bottom three rows, marked as alpha, gamma, beta), one for each joint, and a number of global units to allow for switching on or off walking and for switching between walking forward or backward, plus the units required for sending and receiving information to and from other legs for interleg coordination (Fig 2, brown and ocher units). Further, there is a global input to all joint controllers representing the walking velocity of the hexapod (leftmost white unit – upper part of Fig. 2, below coordination units shown in brown). This unit receives input between 0 mV and 50 mV. There is sensory feedback from the joints in form of a simplified antagonistic sensor structure illustrated by the black squares containing piecewise linear functions (input: angular degrees, alpha pos., beta pos., gamma pos., output: neuronal excitation in mV). This structure projects on an artificial neuron allowing for activation between 0 and 50 mV.

Correspondingly, we simplify the motor output by considering two antagonistic muscles for each joint, the protractor-retractor group for the alpha joint, the levator-depressor group for the beta joint and the flexor-extensor group for the gamma joint. In the simulation, each muscle pair is represented by one motor as explained above.

Next, we will explain how sensory input elements and motor output elements are connected.

##### Control of Beta Joint (bottom part of Fig. 2)

The beta controller represents the most simple case: Each of both antagonistic branches (Fig. 2, levator above, depressor below) is designed as a negative proportional position feedback (NPPF) controller: each branch receives its sensory input (joint position value, between 0 mV and 50 mV) that is subtracted from a given set point (rightmost red units for each branch). ‘Proportional position controller’ means that the negative feedback controller behaves like an elastic spring. The value of the set point depends on the state of the leg (swing or stance; these are channeled from light grey units in upper part of Fig. 2) and is given by a small network called height net (beta controller, blue and red units, for details see below). The error signals (output of red units marked “error”) are then given to the premotor units (PMN, left, units connected via inhibitory neurons) which in turn project to the motor units (leftmost red units, MN). In all joints, the motor output represents a velocity signal which is transformed, via the motor, to a torque signal. To minimize co-contraction, the premotor units of both branches are connected via recurrent lateral inhibition (for details see below).

Behavioral data (Cruse et al. 1993) suggested the interpretation that, during stance movement, body height, i.e. the distance between body and ground, is controlled by NPPF controllers which operate independently in each leg. In the simulation, we apply a simple network for control of body height. Basically, the beta controller, due to its property to produce proportional error deviations, could already serve as a primitive height controller. However, if the gamma joint adopts large angle values differing too much from a vertical position of the tibia, height control based on the beta controller alone would produce inappropriate set point values. Therefore, signals from gamma angle are also exploited in height net to improve height control. This is done following Fink et al. (2014) via presynaptic inhibition to implement a multiplicative dependence (called devisive normalization) via the synaptic input to the central unit of the three rightmost lower red units (Fig 2). Fig S1 illustrates how the set point value of the beta controller depends on gamma angle in a range of 60 to −57 degrees. The range used during normal walking amounts between −9 and −23 degrees.

In the beta controller, when swing mode is started, a unit with bandpass properties (uppermost large green unit, marked HPF) is activated to “disturb” the position controller in such a way that the levator branch is briefly activated leading to a levation of the leg (Schumm and Cruse 2006). As the activation of this unit ceases, the negative position feedback controller pulls the beta joint back to its set position. This “disturbance” signal is applied to agree with the observation that the maximum amplitude of swing movement is constant independent of starting level of the leg (Schumm and Cruse 2006). The signal from this unit may also be given to the extensor branch of the gamma joint controller, which helps to lift the tip of the leg higher up during swing. For very slow velocities this “disturbance” is smaller, leading to short steps type P (see below).

The controller for the beta joint can be used for both forward walking and backward walking.

##### Control of Alpha and Gamma Joint

Alpha controller and gamma controller have to handle forward and backward walking differently. The structure of both alpha controller and gamma controller requires two symmetric branches: Control distinguishes between different behaviors, swing and stance. The mapping from swing and stance to activation of the antagonistic motor neurons is different in forward and backward walking. The activation of the two motor neurons is actually reversed in the case of backward walking with respect to forward walking. Therefore, it is necessary that the input signal for swing as well as that for stance can be given to both motor output units of each joint.

The controllers for swing movements are based in principle on the same structure as the beta controller, i.e. show a NPPF controller. An error signal is calculated from the currently sensed joint position (given by the black squares addressed above) and a set point provided as a target position for the swing movement (set points may vary during negotiation of curves, see Supplement). In the alpha controller and the gamma controller the error signals (two right large green units in the alpha branch and in the gamma branch) are given to the premotor units via an additional parallel feedforward connection (corresponding leftmost small green units). These connections guarantee a minimum constant output value until the error signal is about zero (i.e. until the joint angle has reached its set point). Note that the set point is beyond the position where end of swing is normally reached as swing is terminated when ground contact is sensed.

Stance movement depends on the global input signal for the velocity represented by a global velocity unit (Fig 2, leftmost white unit). This unit can adopt values between 0 mV and 50 mV (in the simulations we use values from 8 mV to 50 mV). For each leg, this global value is transmitted by a special network (“ring net”, Fig 10) that determines the velocity parameter entering the alpha branches and the gamma branches, via the blue units, at its rightmost red units, respectively, but only during stance (see Gabriel and Büschges 2007). These signals depend on the alpha position of the leg under view and the angle characterizing the currently desired walking direction theta (equals zero degrees for walking straight forward). This ring network will be explained below, after briefly explaining how switching between the two behaviors takes place.

##### End of swing, end of stance

Before we turn towards coordination of legs, we want to briefly summarize and explain how the single leg controller controls behavior over a single cycle. In particular, the focus shall be on the transitions between the two different behaviors, stance and swing.

Swing movement is finished at the latest as soon as the swinging leg has reached a given anterior position. If, however, the “torque” signal from the beta joint is higher than a given threshold, the leg is considered under load, and this information is used to stop swing even if the leg has not yet reached the prescribed anterior position of the alpha joint. In this case, the rightmost unit of the group of four light grey units (representing state of stance) is activated by the torque signal and protractor motor neuron is inhibited. This mechanism takes however place only if the leg has reached a given position during swing movement as observed in stick insects (Schumm and Cruse 2006).

Correspondingly, end of stance is determined either by reaching a given leg position in extreme cases or, normally, by earlier signals from any interleg coordination influences. A direct influence of load to suppress swing is not implemented. Instead, stance motivation (via the rightmost turquoise unit) is decreasing versus the end of stance which allows stronger coordination influences to finish stance earlier (Cruse 1985c, Duysens et al 2000). As a specific property to simplify start of swing, the protractor output is inhibited as long as the leg is already in swing state but still under load. In this way, the leg can be lifted in vertical direction and slipping over ground elicited by possible extension of the gamma joint is minimized.

Two types of short steps have been observed (Bläsing and Cruse 2004, Theunissen and Dürr 2013;). The first has been termed type A (for anterior) in the Discussion. Here, we did implement another type of short steps, termed type P (for posterior). During very slow walking, while trying to search a way to cross a gap, stick insects (*Aretaon asperrimus*) may perform swing movements starting from normal PEP but with very small amplitude (Bläsing and Cruse 2004). This behavior could be simulated by neuroWalknet if, for very slow walking velocities, the output of the uppermost large green unit is decreased. Thereby the levator would be lifted to a small degree only resulting in a short swing movement. This type of short step is required to simulate negotiation of tight turns as given in Supplement (Fig 2, two uppermost green units marked “short step P”) for details explaining the box “local(theta)” in Fig 2 see Suppl).

To summarize: The alpha controller and the gamma controller represent a velocity controller during stance and a proportional position controller during swing using the PID velocity controller as a lower level cascade. In the beta joint, proportional position control is applied during swing and during stance.

##### Adaptation of Leg Movement depending on Walking Direction – Ring Net

To allow for an approximately straight trajectory of the leg tip, the contribution of the three leg joints should be controlled accordingly (in the current approach contribution of angle beta is neglected). In addition, curve walking requires a change of the stance trajectory of the leg (Supplement, Fig. S3). To this end, each leg is equipped with a “ring” network (Fig 10) that controls the alpha joint and the gamma joint during stance to provide a leg trajectory that contributes to walking in a direction given by angle theta (e.g. theta = 0 for all legs when walking straight forward). This network consists of five layers each containing 12 units (shown from right to left in Fig. 10A, Fig 10B gives an overview showing the complete ring structure schematically). Functionally, the units of layers 1 – 3 (Fig 10B, green ring) act as binary elements. The first layer (Fig. 10A, rightmost column, delta) is activated by sensory input representing the angle delta (delta = alpha* – theta, Fig 2, box “spatial coding”) with alpha* = (alpha + 50)/3.). Angle alpha is provided by the protractor branch, i.e. the sensor cell representing the alpha angle that increases activation from rear to front. The desired walking direction for each leg is provided by the box “local(theta)” (Fig 2, upper part, below white units). Function local(theta) might be different for different legs depending on the curve radius, i.e., global walking direction. The transformation of rate coded delta to spatial coded delta is not implemented explicitly (Fig 2, box “spatial coding”). Rather, parameters are given for a specific example of curve walking only. Fig 2 illustrates the dependence graphically.

Angle delta is represented by using so-called spatial coding. As 12 units are given for an angular range of 180 degrees, one unit represents an angular range of 15 degrees (Fig. 10B). This means that for alpha = 30 degrees and desired direction theta = 0, i.e. straight forward, for example, in the first layer all units from C1 to C6 are excited (see Fig 10A, green dots).

Layer 4 and layer 5 (Fig 10A) control velocity of alpha joint and gamma joint, respectively. The function of layers 2 and 3 is to produce activations in layers 4 and 5 in such a way that only one unit is activated to represent the position of angle delta: First, in layer 2 only one unit is activated, the one that specifies the current delta angle. In the example depicted in Fig 10A this would refer to unit C6 (green dots). Layer 3 represents an inverted version of layer 2 which is required as we need active inhibition of all units in layers 4 and 5 by the units not activated in layer 2 except the one being activated in layer 2. All units of the third layer are then projected to a forth and a fifth layer in parallel. These units receive input from the unit representing the desired walking velocity (Fig 10A, vel; Fig 2, box “local(theta)) via the fixed weights “0.2* sin(alpha)” for layer 4 and “0.2*cos(alpha)” for layer 5. All 12 units of layer 4 (Fig 10B, red) project to output unit #O2 (currently, unit O1 is not yet connected to protractor, as the latter is required for negotiating tight curves only). Units C81 to C86 of layer 5 (Fig 10B, yellow) project to unit O4, units C87 to C92 (Fig 10B, blue) project to unit O3. Units O1 – O4 represent the contribution of the alpha joint (O2) and the gamma joint (O3, O4), respectively, to produce an appropriate leg movement. Units O1 and O2 in Fig 10A correspond to the rightmost dark blue units addressing the alpha branch (Fig 2, upper) and units O3 and O4 to the corresponding units addressing the gamma branch (Fig 2, lower). As depicted in Fig. 2, the four left dark blue units of the alpha branch (upper) and the gamma branch (lower) respectively receive – via the small dark blue units – this information and are responsible to direct the signals to the appropriate motor units depending on (i) position of the leg, (ii) the current state, swing or stance, and (iii) forward (including curve walking) or backward.

Taken together, the ring net as used here represents an approximation of a vector with the motor output components of alpha joint and gamma joint pointing to the desired walking direction theta (e.g. straight forward, theta = 0). It is an approximation only because (i) the angle values used have a limited resolution (accuracy 15 degrees) and (ii) the contribution of the beta joint is neglected. Nonetheless, the behavioral results obtained in the simulation show that such an approach is sufficiently able to describe the behavior of the insect (the relation of the ring network to a neural structure found in the central complex is addressed in the Discussion).

As a general property, not explicitly required for straight walking, we added a limitation of stance movement not only with respect to the alpha angle, but also to the gamma angle (upper limit = 35 mV, Fig S1). This is required for negotiating curves as, due to the oblique trajectory, the inner front leg would not reach the alpha thresholds before the leg had moved below the body (see Fig S 3). The existence of such limits follows from data given by Dürr and Ebeling (2005) and also by Gruhn et al. (2009).

#### C) Interleg coordination

Behavioral observations in forward walking stick insects have led to five “rules” that may underlie interleg coordination (Dürr et al. 2004). In these experiments various disturbances, essentially prolongation or shortening of swing duration or stance duration, have been applied to animals walking either unrestrained, or supported by a holder on a treadwheel or on slippery surface. Here we deal with rules 1-3 first (all units marked brown, Fig 2; these have been realized in Walknet as well, Schilling et al. 2013a) and later rule 5 (all units marked “ocher”, Fig 2; this has been realized in an earlier version of Walknet, Schilling et al. 2007). We will not test rule 4, which describes the faculty of one leg to use the position of its anterior neighboring leg as a “target”, or set point, for the goal of its swing trajectory. One reason for that is that the size of front legs and hind legs in the robot do not fit to those of the stick insect, making simulation of targeting behavior difficult. Another reason is that it is not known to what extent targeting may be activated or not (see Supplement). We also do not address interesting results showing that coordination between legs is also driven by mechanical coupling between legs (Dallmann et al. 2017) which may support – or replace – rule 1 as a neural mechanism, at least when walking on horizontal substrate.

All rules connect only directly neighboring legs and depend on state (swing, stance) of the sender leg as well as on the state of the receiver leg (see Fig 2, Fig 11). In Fig 2 the connections to and from other legs are depicted by bold dark arrows.

*Rule 1* is assumed to act only between ipsilateral neighboring legs in anterior direction (rear to front). It inhibits start of swing in the anterior leg and is split into two sections (1a, 1b). Influence 1a is active as long as the sender leg is in swing state (brown units, output 1a). Influence 1b represents a temporal delay driven by an integrator (characterized by a recurrent self-activation), the output of which is however inhibited when rule 2i is activated, too (see below).

**Fig. 11.**
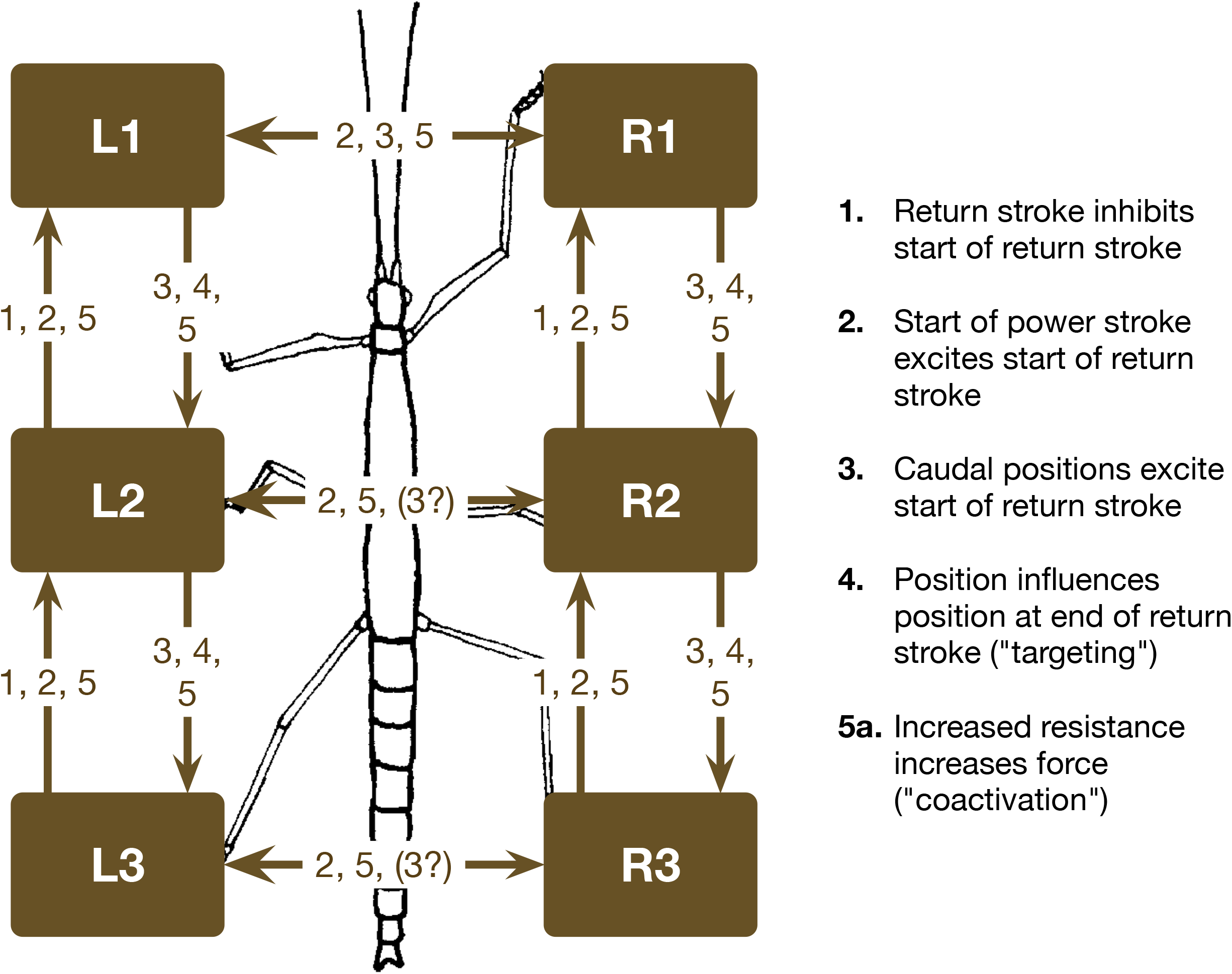
**Single leg controllers (shown in brown) and their connection via coordination rules** (**from Durr** et al. 2004). L1, L2, L3 left front, middle, and hind leg, respectively. R1, R2, and R3 stand for the corresponding right legs. The question mark indicates that there are ambiguous data concerning influence 3.

*Rule 2i* (i stands for ipsilateral) depends on velocity via the rightmost of the three lower brown units inhibiting the output of the integrator. As soon as inhibition of this unit is finished, two serially connected units (marked by “HPF”, together forming a second order high pass filter) activate an output signal that represents rule 2i. This effect may excite swing in the receiver leg, so that inhibition of swing via 1b is directly followed by excitation of swing. In other words, rules 1 and 2i together may trigger swing in the anterior neighbor in such a way that, after a short delay following the end of swing movement of the sender, swing of the receiver leg is started. For high velocities the delay is short, but increases for slower velocities (see Graham 1972).

##### Rule 3i

Complementary to rule 2i there is rule 3i, as it elicits swing, too. However, rule 3i operates ipsilaterally from a sender leg to its posterior neighbor (i.e. from front leg to middle leg, and from middle leg to hind leg). This influence depends on alpha position of the sender leg and switches on (thr_on) or off (thr_off) the sender signal which depends on leg position as well as a threshold function depending linearly on velocity. As both influences, 2i and 3i, superimpose in the case of the middle leg, parameters should ideally be chosen in such a way that 2i being sent from the hind leg starts together with 3i being sent from the front leg so that both influences support each other. However, longer duration of 3i activation may act to move not yet optimally coupled legs nearer to optimal values.

##### Contralateral – rule 2c and 3c

Further, there are two influences 2c and 3c (c for contralateral) that act between contralateral neighbors and act symmetrically in both directions. Influence 2c is dependent on the position of the sender leg via the unit marked “HPF”. Rule 3c depends on position of the sender leg, too, but in this case there is no high pass filter. This means that the effect of 2c decreases with decreasing velocity, in contrast to influence 3c.

##### No Rule 1 contralateral

The effect of rule 1 has first been postulated by Hughes (1952). First indirect experimental evidence pointing to this effect has been found for Neoconocephalus between hind leg and middle leg by Graham (1978) and indirectly for contralateral legs in *Carausius* (Foth and Graham 1983). Direct experimental evidence has however only been revealed among ipsilateral legs (hind leg to middle leg, middle leg to front leg), but not for contralateral legs (Cruse and Epstein 1982). Therefore, rule 1 influences for contralateral legs have not been implemented.

##### Backward Walking

For backward walking, rules 1 and 2i are used in the same way as during forward walking. Rule 2c for backward walking corresponds to rule 2c for forward walking, with a however different position threshold. Rule 3i and 3c are not activated during backward walking.

##### Rule 5

In addition to rules 1-3, rule 5 (5i, 5c) is implemented, too (here we refer only to rule 5a, not to rule 5b, Fig 11). Rule 5 differs fundamentally from the other rules as it depends on the output of the retractor premotor unit of the sender leg (shown as the upper leftmost unit marked in ocher, output 5) and that it excites the retractor premotor unit of the receiver legs while at the same time inhibiting the antagonistic protractor premotor units (for other joints, see below). In other words, these influences support cooperative (in-phase) motor output of legs being coupled. In Fig 2 input via rule 5i is depicted as the upper rightmost unit in ocher. The left neighboring unit receives input 5c. Rule 5 acts between all neighboring legs except both hind legs (Cruse 1985a). The influences are only effective if the retractor of the sender is strongly activated. In other words, during undisturbed walking rule 5 influences are not activated. For the hind leg, we further introduce rule 5ch (leftmost unit of rule 5 input), which shows the same kind of connectivity, but inhibits the retractor and excites the protractor premotor units of the receiving leg, i.e. supports anti-phase coupling, an assumption strongly inspired by results of (Knebel et al. 2017).

Several studies concerning the coupling of CPGs (Büschges et al. 1995, Mantziaris et al. 2017, Knebel et al. 2017, Borgmann 2007, 2009) reported comparable results for recordings from retractor motor neurons (i.e alpha joint) and from depressor neurons (i.e. beta joint). As a consequence, rule 5 has been introduced to both joint controllers. As Büschges et al. (1995) showed that oscillations of alpha joint and beta joints are not coupled with each other, both joints are connected independently via rule 5.

A number of behavioral and neurophysiological studies support the assumption that coordination between ipsilateral legs is stronger than between contralateral legs (for walking, e.g. Cruse 1990, for deafferented preparations e.g. Knebel et al. 2017). To agree with this property, rule 5 connections between ipsilateral legs are coupled by a factor of 0.2, whereas connections between contralateral legs are connected via a factor of 0.1.

### Specific Experimental Settings

The simulations have been applied to simulate a wide variety of different locomotion experiments. In these cases, different velocities were used and, for moderately fast velocity, curve walking was tested. Furthermore, we performed simulations that parallel specific experimental settings: on the one hand, experimental setups in which animals were put on a treadmill where in some cases only a couple of legs were walking. On the other hand, experimental interventions in which the animal was deafferented and as a consequence there was no sensory input from the animal nor movement produced. In the following, we will describe how we simulated these experiments.

#### Setup of Treadmill Experiments

In some experiments (Borgmann et al. 2009; Clarac and Chasserat 1979; Cruse and Saxler 1980) animals were fixed on a holder and walked on a treadmill, whereas other legs were deafferented or were intact but standing on a fixed substrate. We assume that rule 5 influences of legs walking on a treadmill are stronger than leg controllers of a ganglion ‘treated with pilocarpine’.

Simulation of motor output required for a single leg moving a treadmill is not straight forward because the forces to be applied by the animals may vary depending on the inertia and friction of the specific treadmill applied. Even the extremely light-weight treadwheel developed by Graham (1981) still had a two-fold larger inertia compared to a free walking stick insect in the horizontal direction and an eight-fold inertia in vertical direction. Applying an indirect measure, Cruse (1985b) could show that front legs show stronger forces on a treadwheel, but no data are available concerning contribution of alpha or gamma joint. Therefore, in order to reproduce – on a qualitative level – results observed by the experiments of Borgmann et al. (2009) and later by Cruse and Saxler (1980), we use the following approximation. Legs walking on a treadmill are assumed to produce higher rule 5 signals at the beginning of stance, due to higher friction to be compensated (see Cruse 1985a). To this end, during the first half of leg position (alpha joint position sensor > 25mV) the factor of rule 5 is increased from 1 to 15, but, to compensate friction effects, does not increase velocity output. This is necessary because friction cannot be simulated in the robot Hector. In the figures, MN output activation is shown if above a given threshold (see legends of Figs 7–9, S5, S7).

#### CPG Experiments With a Deafferented System

Although the general architecture of neuroWalknet does not rely on CPGs to control normal leg movement (i.e. very fast velocity is given), the network contains elements that under specific conditions may operate as oscillators.

As mentioned above in this section, the antagonistic branches of each joint controller are connected via lateral inhibition to avoid co-contraction. A network consisting of two units connected via lateral inhibition could be made to oscillate, if (a) we deal with backward (recurrent) inhibition rather than forward inhibition, (b) if inhibitory connections are strong enough, (c) if recurrent inhibitory units show adaptive (i.e. high pass filter like) properties, and (d) most important, if both units receive parallel excitation. In our case the latter condition is normally not given during normal walking, as we deal with an antagonistic architecture, where in general only one of both channels is activated. However, if sensors are either deafferented or, in addition, treated with pilocarpine, both channels are activated and therefore oscillations can be elicited (Büschges et al. 1995). This may be the case for different sensor systems. For simplicity, here we address them by black input arrows marked “pilo”. This may reflect campaniform sensilla, but any other mechanosensor may be suited to elicit this function, too. As there are no reports concerning mutual inhibition to exist between motor neurons, these connections are implemented on the premotor level. The fictive load sensors are implemented for each joint but used only to activate both premotor units to allow for simulation of treatment with pilocarpine, i.e., respond to strong activations only. To simulate experiments with deafferented (e.g. autotomized) and/or pilocarpine treated animals, we fix the motor output (joint angle velocity) to zero in all joints, and set all load sensor inputs (“pilo”) to high values (>= 25 mV).

## Supporting information

Video - fw walking, slow, velocity 8

Video - fw walking, slow, velocity 10

Video - fw walking, slow, velocity 15

Video - fw walking, slow, velocity 20

Video - fw walking, velocity 25

Video - fw walking, velocity 30

Video - fw walking, velocity 35

Video - fw walking, velocity 40

Video - fw walking, velocity 45

Video - fw walking, velocity 50

Video - curve walking

Video - running

Video - bw walking, velocity 20

Video - bw walking, velocity 30

Video - bw walking, velocity 40

Video - bw walking, velocity 50

## End Matter

## Author Contributions and Notes

M.S. and H.C. designed and performed research, H.C. and M.S. wrote simulation software and developed concept of the model; M.S. and H.C. wrote the paper.

The authors declare no conflict of interest.

This article contains supporting information online (Videos and Supplemental Material).

## Acknowledgments

The authors thank Thierry Hoinville, Josef Schmitz, Axel Schneider, and Volker Dürr for discussions and helpful comments on the manuscript. This work was supported by the Cluster of Excellence Cognitive Interaction Technology CITEC (EXC 277) at Bielefeld University, which is funded by the German Research Foundation (DFG).

