## Supplement for "Decentralized Control of Insect Walking – a simple neural network explains a wide range of behavioral and neurophysiological results"

The supplement adds, first, further details on the control structure (A – height control); second, provides information on motor activations over one stepping cycle (B); third, provides details for one example of curve walking and results for different velocities (C – Negotiating curves; D – Footfall patterns for different velocities; E - running) and, lastly, discusses design decisions on the controller (F – Selection of coordination influences and differences between species).

### A) Height Control

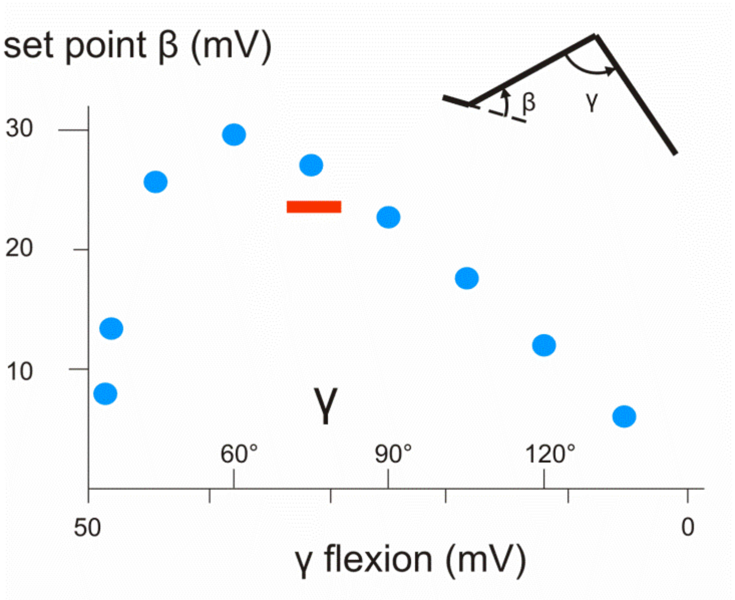


**Fig S1.** Body height depends on angles beta and gamma (the contribution of angle alpha is neglected). The network controlling body height during stance is depicted in Fig 2 (three blue units, lower right) and is actively controlling the set point of the beta angle (see Methods, Control of beta joint). Here it is shown how, during stance, the set point for the beta joint controller depends on the gamma angle (see inset visualizing how both define body height). This function is approximated by presynaptic inhibition (Fig 2, input to the central unit of the three blue units, lower right, as depicted by the blue dots). Abscissa: gamma angle (degrees) and flexion (mV), which is used as input for height control. Ordinate: set point for beta controller during stance (mV), as depicted in the inset. The red bar marks the range of gamma angles used in the simulations during normal walking.

**B) Motor Activation during Stepping Cycle**

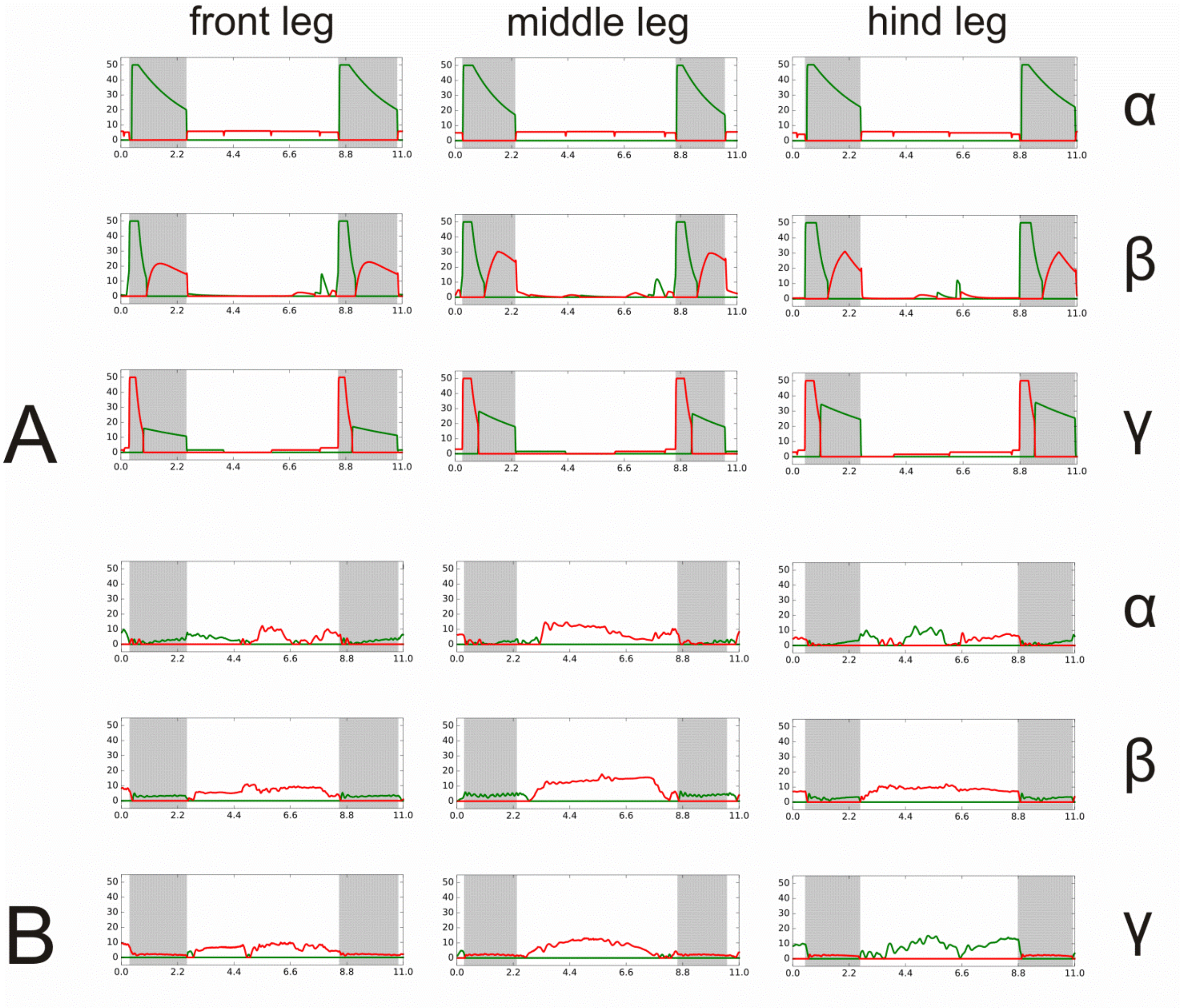


**Fig S2**. Sections of steps for different legs. A) Motor output provided by the six motor neurons (Fig 2, MN). B) Approximation of torque values provided by the six muscles for the same steps as in A (see text). Walking velocity 30 mV. Alpha joint: protractor green, retractor red; beta joint: levator green, depressor red; gamma joint: flexor green, extensor red. Swing state: grey shadows. Abscissa time (s), ordinate (mV).

Fig S2A shows motor output provided by the six motor neurons (Fig 2, MN) and Fig 2SB shows an approximation to torque values provided by the six muscles of the three legs (front leg, middle leg, hind leg) during a complete step cycle. Sensory input required for performing a swing movement or a stance movement is given by the light grey units (Fig 2). These units receive sensory input (position and load) from the own leg and from other legs via interleg coordination influences. Swing state, as represented by activation of the leftmost light grey unit in Fig 2, is highlighted grey in Fig 2. Within each mode, motor units of the different joints are, during swing, controlled by negative feedback controllers, which, depending on the current leg position, may activate one of both antagonists. During swing, a HPF signal acts as a temporally limited disturbance signal at the beginning of swing thereby lifting the leg. Set points are fixed, but are different for different joints. During stance, the motor output of alpha joint antagonists and gamma joint antagonists receive signals via the ring net which depend on the current angle alpha (and the global walking direction). The beta joint is governed by a proportional negative feedback controller the set point of which depends on gamma angle, in order to maintain an about constant body – ground distance (Fig S1). Depending on the somewhat different geometrical position of the legs (and the different position of the COM), the temporal order of antagonist activations is different for the different legs. Note that switches between antagonistic muscles can be found at different temporal moments during the complete cycle, even within swing mode or stance mode. As discussed by Dalmann et al. (2016), motor output activity and torques may operate in opposite direction as is obvious in the second part of gamma joint of the hind leg or the first part of the gamma joint in the middle leg.

Simulation results depicted in Fig S2B may be compared with torques received from stance movements of free walking stick insects (Dahlmann et al. 2016). In all the legs, there is a good agreement with respect to the beta joint, where the depressor (red) provides the torque required to support the body. Qualitative agreement can be observed as well in the gamma joints. The extensor torque dominates the stance of the middle leg and the second part of the front leg in both biological results and simulation. The second part of the hind leg stance is characterized by flexor torque in both simulation and biological experiments, in spite of extensor velocity activation. Results differ in the first part of the front leg stance, where simulation shows extensor torque activation, whereas the biological experiments show flexor torque activation. In the first part of hind leg stance, simulation shows flexor activation whereas the biological experiments show extensor activation. Concerning the alpha joint, there is good agreement with respect to the front leg and the middle leg. The hind leg simulation shows an irregular alternation between protractor and retractor torque, as does the biological data. However, the tendencies are different. In the biological data retractor dominates in the first section of stance, and protractor in the second part, opposite to that found in the simulation.

Taken together, there is good overall agreement for middle leg and for the beta joint of front leg and hind leg, but no clear match in alpha joint and gamma joint of the front leg and the hind leg, where the biological data do not show a very clear picture either. Note that a perfect fit is not to be expected due to the difference of leg geometry and, importantly, different position of center of mass in both cases (see Methods).

### C) Results – Negotiating Curves

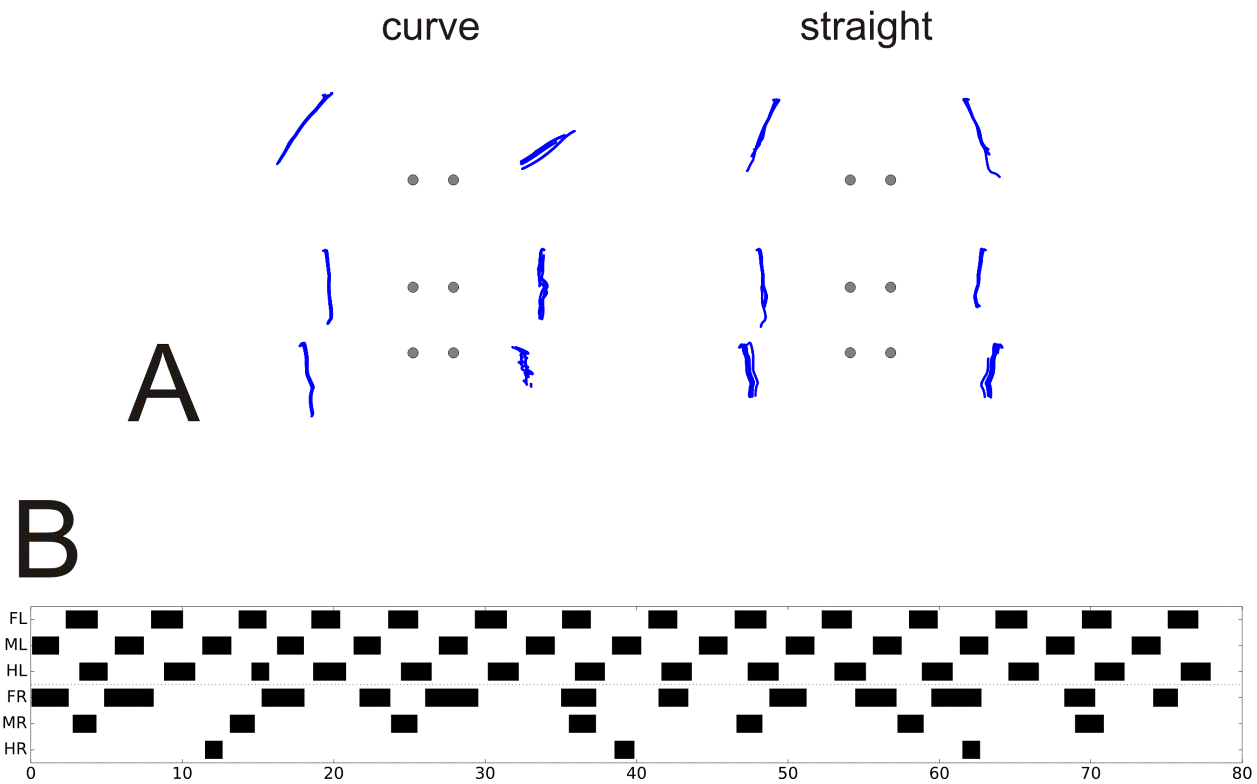


Fig S 3. Negotiation of Curves. A) Trajectory of leg end points during stance in curve walking (see left) with respect to a body centered frame of reference, plotted over time window 10 s – 100 s. On the right, trajectories of leg end points during straight walking is shown for comparison. Dots mark position of leg basis. Front legs up. B) Footfall pattern. Global velocity 45 mV, theta 75 deg. Ordinate: legs as in Fig 3, abscissa: time (s).

Only few data are available concerning negotiation of curves by insects (Rosano and Webb (2007), Gruhn et al. (2009), Dürr and Ebeling (2005), Dürr (2005), Cruse et al (2009), Frantsevich and Cruse (2005). Dürr and Ebeling (2005) show an experiment where simultaneous recordings of footfall patterns, direction of turn as well as AEP and PEP of all six legs have been collected. We focus on this example as a proof of concept as it is the only case we are aware of where all these data are given for a turning insect. The insect walked tethered on a holder above a sphere and curve walking was elicited visually by a horizontally moving pattern of vertical stripes. In these experiments the outer legs show a pattern quite similar to that observed in normally forward walking animals when performing a tripod pattern, whereas the inner legs strongly deviate from that pattern. The inner front leg moves in anti-phase to the outer front legs, as is the case in normal forward walking. However, in this example, the inner middle leg shows an about two times longer step period compared to the inner front leg or all the outer legs. The period of the inner hind legs was even longer and more irregular and showed shorter swing duration compared to straight walking. As swing movements are very short and may show small amplitudes, not each swing might have been recognized. At least the front legs show a shift of the AEP in the direction of the desired turning direction (angle theta) (Dürr and Ebeling (2005), Dürr (2005), Gruhn et al. (2009), Rosano and Webb (2007)). Corresponding deviations from normal forward walking are also observed in the other legs. However, as has been argued on the basis of combined experimental and simulation studies, Rosano and Webb (2007) conclude that the contribution of front legs is sufficient to explain turning behavior of intact stick insects.

To test this hypothesis with neuroWalknet, we kept coordination rules as in normal forward walking (although Dürr (2005) found decreasing coupling strength between inner legs), but implemented an expansion symbolized by box "local(theta)" in Fig. 2, the function of which will be explained as follows. Simulation of negotiation of curves was realized by application of (i) different set points for swing, (ii) different local walking directions, as well as (iii) different velocity signals for different legs instead of using one global velocity for all legs.

How are directions of front leg trajectories influenced? For control of swing movement, the beta controller is used as during forward walking, but the set points of the front legs are now specified correspondingly. The set points for the AEP of alpha joint and gamma joint appear to depend on angle theta. As detailed data are available for one specific case only, we could not aim for a general solution, but, in order to approximate data of Dürr and Ebeling (2005), chose plausible set point values for the inner front leg (alpha = 20 mV, gamma = 20 mV) and for the outer front leg (alpha = 50 mV, gamma = 45 mV). During stance, the contribution of alpha joints and gamma joints are determined by the ring network (see Methods) as in forward walking, i.e. by angles delta = alpha* – theta. In the case of curve walking, theta deviates from zero. Following Rosano and Webb (2007), for the inner front leg theta is set to point to the desired walking direction, e.g. the direction of a goal to be approached. For the specific case, we set theta = 5 (corresponding to an angular range of 75 (± 7.5) deg). As there is no detailed information given concerning the outer leg, as a first approximation to data of Dürr and Ebeling (2005) and Rosano and Webb (2007), we reduced theta for the outer front leg by a factor of 0.25.

How are the specific local velocities determined? To approach the specific data of Dürr and Ebeling (2005) we adapted global velocity (in the example 45 mV) by factor of 1 for all three outer legs and factors of 0.75, 0.35 and 0.1 for the inner front leg, middle led and hind leg, respectively. Again, as only this data set is available, we did not attempt to find a general solution for all possible turning radii, but focused on this special case.

For simulation, all these parameters (position of AEP, theta, local stance velocity) have been stored in "local(theta)" (Fig 2) and are given to the swing set points of both front legs for swing and the local velocities as well as the local delta values to the ring net of each of the six legs, correspondingly, for stance. Due to the very slow velocity of the inner hind leg, swing movements are performed as "short steps", type P (see Methods and Fig 2, "short steps").

Results are shown in Fig S 3A and Fig S 3B in top-down view (duration 100 s). A 90 degree turn required about 30 s, the radius of the circle amounts for about 0.8 body length. The footfall patterns (Fig S3B) show qualitative similarity to those found by Dürr and Ebeling (2005, their Fig 2B). The stance trajectories (leg end point trajectories during stance in top view, Fig S3A, left) are similar, too, but there are also clear differences. The amplitude of the inner front leg is smaller compared to those of the animals (Dürr and Ebeling 2005, their Fig. 2 Aii and Fig 6B). This is mainly due to smaller leg lengths of the robot legs. Further, in Fig S3A, left, the trajectories of the inner middle leg are parallel to the axis along the body, but are more inclined in the insect (for comparison, Fig 3A right, shows trajectories of straight walking, similar to results of Dürr and Ebeling 2005). Such deviations are even stronger when very tight turns are taken (Cruse et al. 2009, Frantsevich and Cruse 2005). This effect is not observed in the simulation because we did not introduce changes of the AEP set points of the middle leg as has been done for the front legs. However, this difference may also be due to the fact that the robot legs may slip during turning which is not the case for insects. Note that changing walking direction as simulated here may happen to start at any point during stance corresponding to results of Dürr and Ebeling (2005) or during swing (Schütz and Dürr 2011).

More irregular steps of the inner hind leg observed in the animals compared to those seen in simulation may be due to swing movements not being recognized in the experiment because the vertical amplitude is small and the leg may not leave the ground due to the current load distribution of the specific leg configuration. Therefore, simulation results recording neuronal activity (Fig S 3) could appear as more regular than the behavioral results (Dürr and Ebeling, 2005, their Fig 2 Aii).

Tests with varying parameters showed that application of different local velocities was indeed necessary to elicit this kind of curve walking. Changing only the AEP of front legs was not sufficient.

Taken together, simulation of this specific case shows good agreement with the behavioral results and might therefore be considered as a proof of concept for control of curve walking. However, more biological data would be required to expand the simulation for being able to describe turning on a more general level.

**D) Results – Forward Walking, Variations of Velocities**

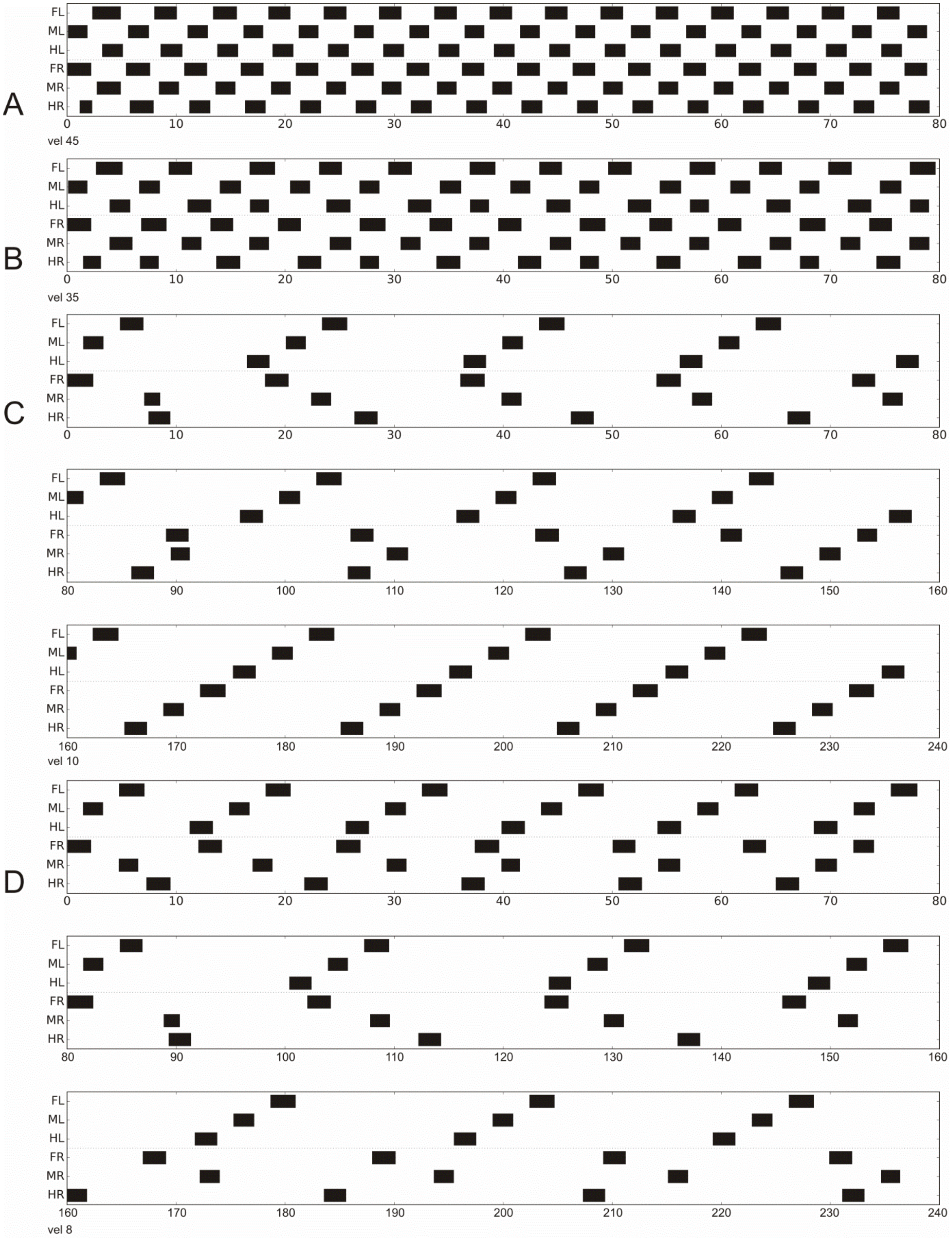


**Fig S4.** More footfall pattern for forward walking, A) vel = 45 mV, B) vel = 35 mV. C), D) as in Fig 3, but now transients are shown, when starting with the usual leg configuration. C) vel = 10 mV, for 240 s, D) vel = 8 mV, for 240 s, for stable pattern see Fig 3H. Ordinate: legs as in Fig 3, abscissa: time (s).

A

B

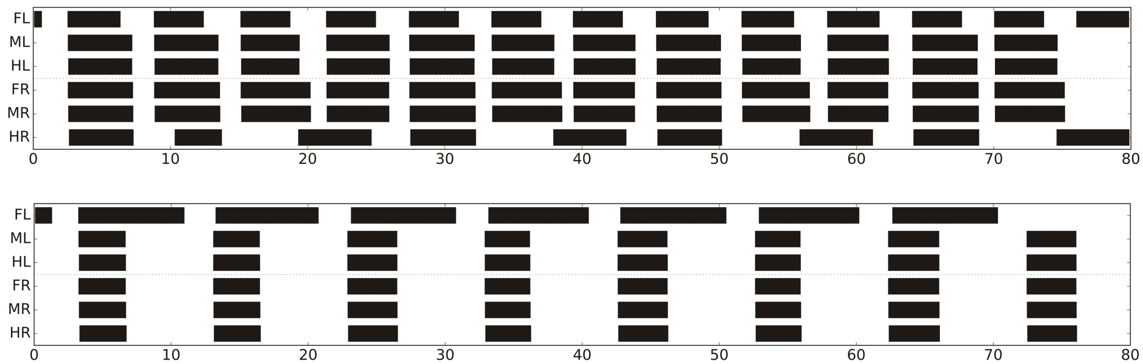


**Fig S5.** Simulation of Borgmann et al. (2009) as in Fig 8, but using different velocities (upper panel, A) vel = 40 mV, lower panel, B) vel = 20 mV). Ordinate: legs as in Fig 3, abscissa: time (s).

**E) Running**

While neuroWalknet can serve as a hypothesis describing the control of hexapod walking covering very slow ("pentapod gait"), medium ("tetrapod gait") and fast ("tripod gait") walking, there are basic open questions concerning control of very fast walking. How might a controller allowing for very fast walking have been developed? In-phase coupling of neighboring hemiganglia, i.e. rule 5-like connections which is found in similar ways in swimming and in flying arthropods, may represent an evolutionary early property. In walking insects, this faculty appears to be only used when the legs are under high load (Cruse 1985a), but not during "normal" walking anymore.

The faculty of very fast walking or running (beyond step frequencies of about 5 steps/s) as observed in cockroaches may represent the result of a subsequent evolutionary development, following the constraint that running does not provide enough time required for sensory feedback. As addressed in the Introduction and the Discussion, for very fast walking ("Run") the sensory delay does not enable control of walking based on sensory feedback beyond a given velocity. For insects of the size of cockroaches as Periplaneta americana, or Blaberus discoidalis this limit is estimated to exist at about 5 - 7 steps /s (Sponberg and Full, 2008, Delcomyn, 1991, Zill and Moran, 1981). Therefore, our network being based on sensory feedback cannot simply be sped up to higher velocities, if the biological conditions should be addressed. The generally proposed solution to this problem is to use CPGs for controlling the rhythmic movements. In the following we show how, using a minimal expansion of neuroWalknet, properties of CPGs might arise and how the two basic problems, intraleg coordination and interleg coordination, might be solved.

First, we introduced a motivation unit (see Conclusions) that when activated stabilizes the network in state "Run". Introduction of such a separate state for running may be justified because specific ("fast") muscles are activated only when step frequency goes beyond the limit of 5 steps/s (Watson and Ritzmann 1998b). The WTA networks used in slow walking to avoid cocontraction (Fig 2, PMN) are, in state "Run", used to control the motor output for fast walking. Second, motivation unit "Run" activates a small network ("SRP") that connects the joints within a leg. Third, unit "Run" activates a network similar to the rule 5 connections allowing for a stable tripod-like gait.

Concerning the CPGs, we simply use the WTA networks given (Fig 2, red units, PMN). These are now actuated by an input from the motivation unit "Run" (to keep Fig 2 as simple as possible we did not depict this unit) instead of the pilocarpine activation (input "pilo) that was exciting the sensory nerves in deafferented preparations. In addition, the appropriate oscillating frequency and the temporal ratio of the antagonist activity could be controlled by influencing the relaxation time constant of the inhibitory units of the corresponding WTA net (red units, PMN, Fig 2). This may be realized by using input from the unit "Run" to threshold the excitation of the inhibitory units of the WTA nets. However, as a proof of concept we will deal with one velocity value only. Therefore, we chose values that provide a ratio of duration of about 1:3 for levator – depressor, and for extensor – flexor pairs and of about 2:2 for the protractor – retractor pair.

To couple the three joints of a leg, we introduce a small network termed spontaneous recurrent patterns (SRP)-net, following results of Büschges et al. (1995). These authors observed in deafferented stick insects an often appearing temporal relation between a switch from levator activity to depressor activity being accompanied by a stop of extensor activity, which was followed, after a short period, by a switch from protractor to retractor activity. We interpret this observation in such a way that connections exist from depressor to flexor, to protractor and, delayed, to retractor as well as from levator to protractor and extensor. This hypothetical network is depicted by a triangular pink box in Fig 2. The complete network is given in Fig S6. Functionally, this network activates levator, protractor and extensor at a moment corresponding to PEP in slow walking, and activates depressor at the upper extreme position during swing, and retractor at a moment corresponding to AEP in slow walking. Note that this is different to earlier approaches that introduced intraleg coupling using sensory feedback (e.g. Büschges 2012, Tóth and Daun 2019).

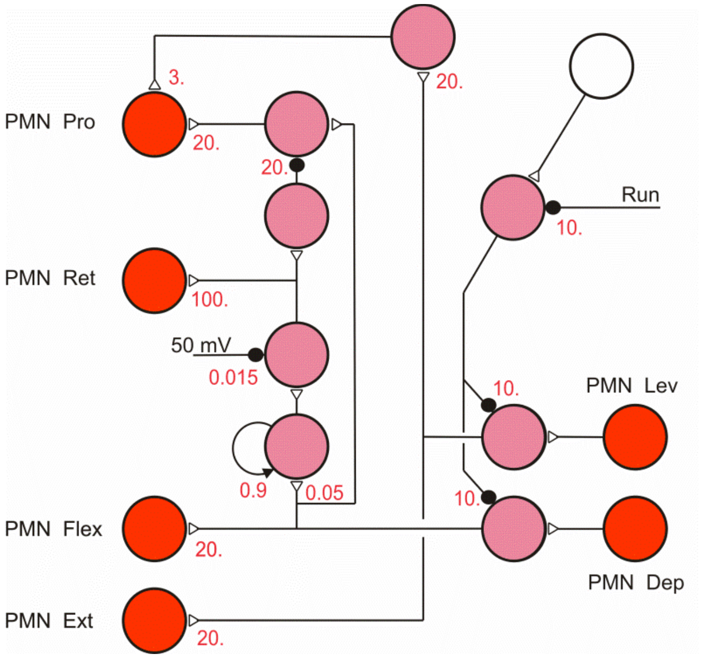


**Fig S6.** SRP-net, used for intraleg coordination during very fast walking (see Fig 2, pink triangle). PMN: premotor units (red), alpha joint: protractor, retractor; beta joint: levator, depressor; gamma joint: flexor, extensor, as depicted in Fig 2. White unit is continuously activated (50 mV).

To address the question concerning interleg coupling, we applied a network (Fig 2, light yellow units) very similar to that used for rule 5, but introduce, based on results of Pearson and Iles (1973), inhibitory influences from a levator unit to the levators of the directly neighboring legs instead of positive feedback as used in the case of rule 5. These 'Pearson-rule' connections lead to a phase distribution in such a way that a levator burst is either directly followed or directly preceded by the levator burst of its neighboring leg (Pearson and Iles, 1973, their Fig 4).

In the simulation, all sensory inputs are suppressed by means of the motivation unit "Run" to simulate the situation where appropriate sensory input is not available. Note that in this simulation interleg coordination relies on mutual inhibition between all directly neighboring legs, whereas for intraleg coordination a feedforward net is used where the levator – depressor CPG controls the other joints. The WTA controlling swing and stance (light grey units, Fig 2) are not activated in this mode as they depend on sensory input.

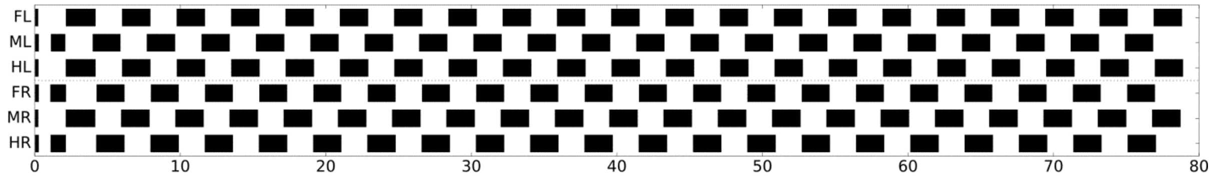


**Fig S7.** Running is illustrated by activity of protractor output (> 0 mV), therefore looking similar to footfall pattern as shown for earlier slow, middle and fast walking examples. Ordinate: legs as in Fig 3, abscissa: time (s).

For testing a solution using minimal feedback we did not influence the CPGs as such, but introduced one position threshold for each joint in a way that the motor output of the joint was set to zero velocity if this threshold has been reached. Application of an at least one-sided threshold allows stable walking patterns, a tripod pattern or possibly some kind of mixture between tripod and tetrapod pattern (see Weihmann et al 2017), resulting from the limitations given by the Pearson rule. An example is given in Fig S7 (+ video) showing a tripod-like case. Note that this pattern (Fig S7) shows activation of protractor output and is therefore similar, but not identical to the footfall patterns showed in earlier figures (showing activation of swing state). We tested different starting positions, but did not perform more detailed investigation concerning the stability of the pattern by using systematic disturbances as has been done for walking.

If only the extreme mechanical limits (i.e. position 0 mV and 50 mV) are given as a threshold, but velocity output to the motors is not switched off, stable patterns can still be observed when focusing on the temporal pattern of the motor outputs. However, to reach cyclic movements of the legs themselves, one has to compensate for possible drift of leg positions. The thresholds required could be realized through nonlinear properties of the muscles for example (Kukillaya and Holmes 2009).

In deafferented cockroaches that were treated with pilocarpine, an anti-phase, tripod-like pattern has been observed (Fuchs et al. 2011). This is in contrast to in-phase coupling found in deafferented locusts, stick insects, rock lobster and crayfish as mentioned earlier. Correspondingly, experiments with intact walking legs and deafferented neighboring legs as performed by Borgmann et al. (2009) with stick insects and by Clarac and Chasserat (1979) with crayfish showed in-phase coupling, whereas cockroaches revealed an anti-phase coupling in this situation. These results (Fuchs et al., 2011) could be simulated, too, if we instead of rule 5 now assume that 'Pearson-rule' network is activated (not shown). Interestingly, a change from in-phase coupling to anti-phase coupling between neighboring legs has been observed in Manduca sexta during metamorphosis (Johnston and Levine 2002), which may be interpreted as ontogenetic recapitulation of a phylogenetic development.

Taken together, we suggest that running may have been developed beyond the ability of slow and fast walking by minor changes of already existing neural systems. For example, CPG properties are gained by common input to both PMNs controlling a joint, whereas the "Pearson-rule" network may have been evolved through a minor change of the rule 5 connectivity. The question whether rhythmic movement is controlled via "minimal neural feedback" or specific nonlinear properties of muscles ("preflexes") that allow for self-stabilization (e.g. Koditscheck et al. 2006, Jindrich and Full 2002) is still open. Near the upper end of the speed limit, different species appear to have found evolutionary solutions that seem to differ in detail (see Weihmann et al. (2015) for a detailed discussion).

### F) Discussion: Differences between species concerning interleg coordination and chosen coordination influences

As to be expected, in spite of similar basic results observed in different species mentioned, there are some differences, too. In rock lobster, the ipsilateral in-phase influence from a walking leg onto a standing leg has only been observed to act in rearward direction (Clarac and Chasserat 1979), whereas in intact stick insects rule 5 influences are provided to both anterior and posterior neighboring legs. In Borgmann et al. (2009) an anterior influence from a walking middle leg to a deafferented front leg has not been found. Mantziaris et al. (2017) found no or weak contralateral coupling among front legs in Carausius m., in contrast to locusts (legs deafferented, Knebel et al. 2017) and stick insects walking on a treadmill (Cruse 1985a, Cruse 1980). These differences may be species specific or may depend on specific methodological properties, for example the concentration of pilocarpine or different friction of the treadmills.

A more general difference appears to exist between coordination rules found in insects (Carausius morosus – Dürr et al. 2004) and crustaceans (crayfish Astacus leptodactylus – Cruse and Müller 1986, Müller and Cruse 1991). In the latter case coordination is stabilized by variation of swing duration instead of duration of stance. This difference has been assumed to result from basic differences concerning the environmental conditions (Cruse 1990).

#### *Coordination Influence 4: Targeting of leg movements*

Targeting is not addressed because it appears to be switched on and off depending on the current walking mode. Targeting seems to be switched off during negotiating of curves, but possibly switched on between front legs during tight turns and may be switched on between middle and hind leg during very tight turning (personal observations). Therefore, targeting behavior appears to be more complex than just operating from hind to middle legs or from middle to front leg as assumed to date. As Schilling et al. (2012) have shown that an internal body model is well suited to support curve walking, one may speculate that the variability of targeting behavior might require some kind of body model instead of a number of separated modules. Data that support this view are given by Dürr and Schilling (2018).

### Simulator and Controller Code

Simulation and code are open sourced: The simulator consists of two parts, one for the neuronal controller and one for the body of the hexapod robot Hector, which exists as a hardware version and as a dynamic simulation (implementations are publicly available: dynamical simulation environment is realized in C++ and based on the Open Dynamics Engine library, for code base of simulator see https://github.com/malteschilling/hector ; the neuroWalknet controller has been implemented in python (version 3), for python code base see https://github.com/hcruse/neuro_walknet ).

### Overview Supplemental Videos

Simulation runs are shown in supplemental videos.

Forward walking at different velocities:

- NeuralWN_vel15.mp4 – forward walking, velocity neuron set to 15 mV
- NeuralWN_vel20.mp4 – forward walking, velocity neuron set to 20 mV

- NeuralWN_vel25.mp4 – forward walking, velocity neuron set to 25 mV
- NeuralWN_vel30.mp4 – forward walking, velocity neuron set to 30 mV
- NeuralWN_vel35.mp4 – forward walking, velocity neuron set to 35 mV
- NeuralWN_vel40.mp4 – forward walking, velocity neuron set to 40 mV
- NeuralWN_vel45.mp4 – forward walking, velocity neuron set to 45 mV
- NeuralWN_vel50.mp4 – forward walking, velocity neuron set to 50 mV
- NeuralWN_run.mp4 – driven at high velocity includes CPG activation

Curve Walking:

- NeuralWN_curve_walking.mp4

Backward walking at different velocities:

- NeuralWN_bw20.mp4 – backward walking, velocity neuron set to 20 mV
- NeuralWN_bw30.mp4 – backward walking, velocity neuron set to 30 mV
- NeuralWN_bw40.mp4 – backward walking, velocity neuron set to 40 mV
- NeuralWN_bw50.mp4 – backward walking, velocity neuron set to 20 mV
